# Eurasian back-migrations into Northeast Africa was a complex and multifaceted process

**DOI:** 10.1101/2022.08.27.505526

**Authors:** Rickard Hammarén, Steven T. Goldstein, Carina M. Schlebusch

## Abstract

Recent studies have identified Northeast Africa as an important area for human movements during the Holocene. Eurasian populations have moved back into Northeastern Africa and contributed to the genetic composition of its people. By gathering the largest reference dataset to date of Northeast, North, and East African as well as Middle Eastern populations, we give new depth to our knowledge of Northeast African demographic history. By employing local ancestry methods, we isolated the Non-African parts of modern-day Northeast African genomes and identified the best putative source populations. Egyptians and Sudanese Copts bore most similarities to Levantine populations whilst other populations in the region generally had predominantly genetic contributions from Arabian peninsula rather than Levantine populations for their Non-African genetic component. We also date admixture events and investigated which factors influenced the date of admixture and find that major linguistic families were associated with the date of Eurasian admixture. Taken as a whole we detect complex patterns of admixture and diverse origins of Eurasian admixture in Northeast African populations of today.

## Introduction

Northeast Africa has undeniably been a key region in human evolutionary history. The out of Africa migrations need to have passed through, if not originated in the region. East Africa is also home to some of the most important fossil evidence for human evolution from early bipedal species such as Australopithecus afarensis [Johanson and Edey, 1990], to the emergence of of early anatomically modern humans such as the Omo fossils [McDougall et al., 2005, Vidal et al., 2022].

Numerous overlapping migrations of farmers and herders over the last several thousand years have also played a critical role in reshaping the current socio-economic and linguistic diversity of the region. It is clear that back migrations into Northeast Africa have had a major impact on the genetic ancestries of the peoples in the region today [Pagani et al., 2012, Prendergast et al., 2019, Molinaro et al., 2019]. Ethiopian populations for instance harbor a large proportion of “non-African” ancestry, as high as, *∼* 40% in some groups – see for instance the Amhara in Figure 1c in Gurdasani et al. [2015]. What is clear is that some current-day Northeast Africans can trace much of their ancestry from other sources than the original hunter-gatherers in the region, such as the Mota individual, an Ethiopian male who lived around 4500 years ago [Gallego Llorente et al., 2015, 2016]. It is also clear that these back migrations into Africa have been ongoing for a long time period. For North Africa, seven individuals from Morocco that had a high affinity to Middle Eastern populations, dated to 15 000 years ago, suggesting the possibility that similar deep-in-time admixtures might have occurred in other parts of Africa [van de Loosdrecht et al., 2018].

**Figure 1:**
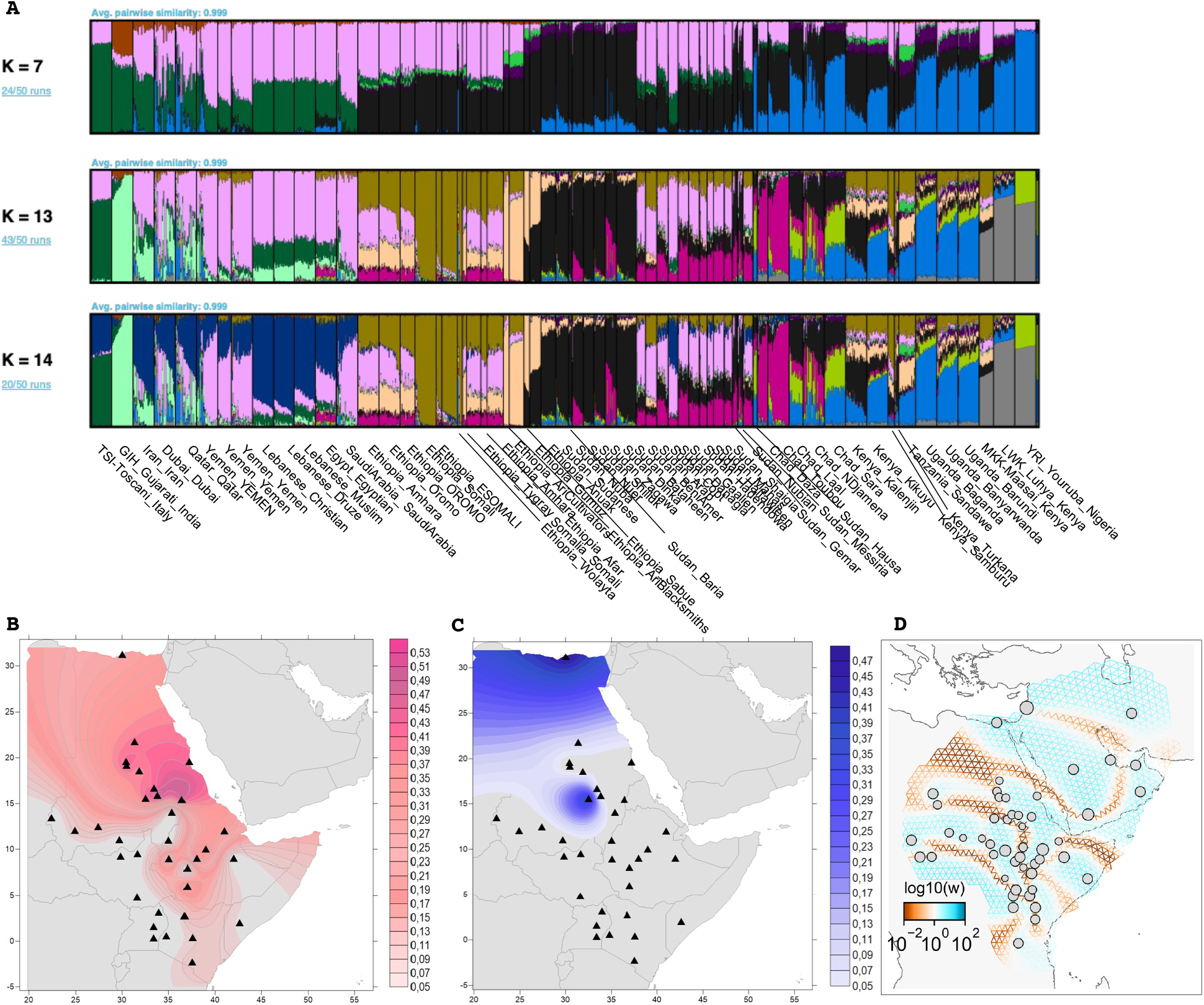
A) ADMIXTURE results for K=7, 13 & 14 visualized using PONG. B) Kriging plot of the distribution of the pink component at K = 14 (maximised in Yemen) in A on the East Africa populations. C) Kriging plot of the distribution of the dark blue component (Lebanese) from K = 14. D) FEEMS plot of inferred patterns of migration rates for the lowest Cross-validation lambda.

In 2017, several ancient genomes were sequenced in an attempt to uncover the demographic patterns in African prehistory [Skoglund et al., 2017]. The study contained data from 16 ancient African individuals from 8 100 – 400 BP. They found that ancient East African hunter-gatherers form a cline of ancestry with modern-day southern African hunter-gatherer (San) groups. This indicates that in the past, hunter-gatherers with a gradient of shared ancestry, ranged from eastern to southern Africa. The fact that these hunter-gatherer groups existed until the relatively recent past allows for the possibility of interactions between them and later pastoralist and agricultural groups in East Africa. This data was later re-analyzed with several new individuals, particularly from East Africa [Prendergast et al., 2019]. They proposed a four-stage model where initially Sudanese Nilotic speakers admixed with groups with Eurasian ancestry (either from Northern Africa or the Levant) within Northeast Africa. In step two, the descendants of these groups migrated to East Africa reaching the Lake Turkana by around 5 000 – 4 000 BP and central Tanzania by around 3000 BP and mixed with local hunter-gatherer groups throughout this process [Prendergast et al., 2019]. The first signs of pastoralism in East Africa coincide with these events. Thirdly the second wave of Sudanese-related groups migrated into the area and contributed to the pastoral iron age populations. Lastly, West African ancestry (genetically similar to Bantu speakers) appeared alongside the advent of crop farming in the region. These findings were then yet again revised in 2020 [Wang et al., 2020]. By analyzing 20 additional ancient individuals, additional resolution was given to the picture and several new patterns emerged. Mainly, Wang et al. [2020] propose that the pastoralists probably arrived in East Africa in multiple waves from several different locations, or that severe population structure was present (distinguishing between the two was not possible). Both Wang et al. [2020] and Prendergast et al. [2019] conclude that there was no single event of hunter-gatherer and herder introgression, neither in space nor in time. Instead a complex “moving frontier” is proposed with diverse patterns of interactions along the contact zones between hunter-gatherer and herder groups.

In the last decade, several genetic studies on modern-day populations have focused on the genetic demographic history of Ethiopia and found patterns of linguistic stratification within Ethiopian populations, i.e. populations within the same language family are more similar to each other than to populations belonging to other language families [Pagani et al., 2012] [Ĺopez et al., 2021], [Hellenthal et al., 2021]. It is less clear if this patterns holds true in Northeast Africa as a whole, as Hollfelder et al. [2017] found a stronger association between geography and genetics than between genetics and linguistic family. By studying modern-day genetic variation, Molinaro et al. [2019] investigated the non-African part of Ethiopian populations and were able to conclude that there has been a Eurasian introgression, likely coming from Levantine, rather than Arabian populations. This event was estimated to have occurred around 3 000 years ago.

By leveraging one of the largest datasets of Northeast African populations to date, we aim to add resolution to Eurasian admixture in Northeast African populations. Specifically, we aim to improve the estimation of the best proxies for the origin of Eurasian admixture in modern-day Northeast African populations by using more Northeast African and Middle Eastern, and Eurasian reference populations. In this study, we follow the approach of [Pagani et al., 2012], [Molinaro et al., 2019] & [Vicente and Schlebusch, 2020] in that we employ local ancestry methods to identify the Eurasian fragments of East African genomes and extract those segments from the surrounding genomes, a process referred to here as ancestry-deconvolution. We then identify the current-day populations that best match those segments. We also date the events to get a better understanding of historic and prehistoric movements in the region. Using the information of possible sources for admixture and dating of these, we construct a model representative of the population history in the region. Overall we find a complex history of Eurasian admixture in Northeastern Africa, related to the spread of languages, the Muslim conquest and trade routes along the Red Sea.

## Materials and Methods

### Genotyping QC

Genotyping data was gathered from previously published studies [Fernandes et al., 2021, Pagani et al., 2012, Gurdasani et al., 2015, The 1000 Genomes Project Consortium, 2015, Rodriguez-Flores et al., 2016, Montinaro et al., 2017, Triska et al., 2015, Patin et al., 2017, Schlebusch et al., 2012, Fortes-Lima et al., 2017, Pagani et al., 2015, Scheinfeldt et al., 2019]. See the table in Supplementary File 1 for a list of populations included in this study, their original population, language classification and geographic coordinates. The geographic sampling information is displayed in Supplementary Figure 1. For this study only autosomal chromosomes were investigated. PLINK v1.90b4.9 [Purcell et al., 2007] was used for data handling and processing. Before data merging, each dataset was quality controlled which entailed 1) removing related individuals using KING [Manichaikul et al., 2010], the first individual within each pair of second-degree relatives or closer was removed 2) SNPs with less than 1% genotyping rate was excluded (`plink--geno` 0.01) 3) C/G and A/T SNPs were removed 4) individuals with at least 10% missingness was removed (`plink--mind` 0.01) 5) potential genotyping errors were removed (`plink--hwe` 0.0000001) 6) lastly only overlapping SNPs between the datasets were kept.

Before analysis that could be adversely affected by linkage disequilibrium (ADMIXTURE and PCA) SNPs in LD were filtered out using `plink--indep-pairwise` 50 10 0.1.

The data from [Pagani et al., 2012] was converted from hg18 to hg19 using the liftOver tool from UCSC (https://genome.ucsc.edu/cgi-bin/hgLiftOver).

As the number of individuals in each population varied quite substantially, from only a few individuals to around a hundred for other populations, we randomly sub-sampled all populations down to 30 individuals. This was done to reduce the effect that sample size can have on the demographic inference.

### Metadata

Geographic information about the populations was gathered from the original publications in the following fashion, 1) directly from text or supplementary tables 2) if no coordinates were provided then they were interfered from the map of sampling locations, 3) if no map or coordinates were supplied, then a point in the middle of the respective country was chosen this was was the case for three publications [Rodriguez-Flores et al., 2016, Pagani et al., 2015, Fernandes et al., 2021]. Regarding language classification, we followed a similar approach as for geographic data, namely that information/classification was used if available in the original publication. The Egyptians from [Pagani et al., 2015] and the Qatari from [Rodriguez-Flores et al., 2016] were both assumed to be Arabic speakers and thus classified as Semitic. The Niger-Kordofanian classification used in [Gurdasani et al., 2015] was changed to Niger-Congo, to better align with the other datasets. For visualization purposes the Semitic speakers on the African continent were given their own label (African Semitic) and their own colour. This distinction was only made to better distinguish between the investigated populations (targets) and Middle Eastern populations used as references. This distinction is thus based solely on geography and is not supported by any linguistic deductions. For a detailed classification of all linguistic groupings used, see Supplementary File 1.

### Population structure inferences

Unsupervised population structure inference analysis for K=2 to K=15 was performed with ADMIXTURE [Alexander and Lange, 2011] version 1.3.0 for K=2 to K=15 using a random seed each time, and repeated 50 times. PONG version 1.5 [Behr et al., 2016] was used to visualize the results and find the major mode and pairwise similarity within the major modes. Principal component analysis (PCA) was performed using FlashPCA version 2.0 [Abraham et al., 2017]. For the PCA plots PC refers to Principal Component, with each value in the PCA plots representing the projection of the data on the eigenvectors, scaled by the eigenvalues. Uniform Manifold Approximation and Projection for Dimension Reduction (UMAP) was performed on the genotypes directly using the umap-learn python library version 0.5.2.

### Patterns of migration rates

The migration rate over the sampling area was investigated using FEEMS [Marcus et al., 2021]. A grid of coordinates covering North-Eastern Africa and most of the Middle East was generated. Cross validation was performed and the lambda with the lowest cross validation value was used to generate the final plot.

### Phasing

Phasing was carried out out using SHAPEIT version 2.r837 [O’Connell et al., 2014] with the 1000 genomes phase 3 reference genomes [The 1000 Genomes Project Consortium, 2015] and options --`states` 500 --`main` 20 --`burn` 10 --`prune` 10. Misaligned sites between the reference dataset and panel were excluded.

### Local ancestry estimation

MOSAIC version 1.3.7 was compiled and ran under R version 4.0.0 [Salter-Townshend and Myers, 2019] to perform local admixture inference, admixture dating as well as ancestry deconvolution. To minimize the potential bias of different sample sizes between investigated target populations, and sources the number of individuals investigated for each population was downsampled to ten individuals. The ancestry deconvolution was performed by running MOSAIC, using the specified resources (see Results for specific scenarios) and then looking at the constructed ancestries that MOSAIC infers from the provided sources. The constructed ancestry in MOSAIC was then compared to the source populations and F_st_ was used to evaluate which one of the source ancestries it most closely resembled. If one of the ancestries shows the most genetic similarity to a Eurasian source then the analysis continued for that ancestry. Thus only samples/targets that mosaic found could be explained by at least one Eurasian ancestry source was ancestry de-convoluted. Segments of each individual’s genomes that were assigned to the Eurasian ancestry with a probability of 80% or more by MOSAIC were saved and the remainder of the genome was set as missing. Admixture dating was extracted from MOSAIC’s co-ancestry curves for the Eurasian like ancestry.

### Outgrup F_3_

Outgroup F_3_ were performed using qp3Pop from `ADMIXTOOLS 2` version 2.0 [Maier and Patterson, Patterson et al., 2012]. The San population Ju’hoansi was used as the outgroup and the extracted ancestry fragments of each target population was tested against populations from the Eurasian reference dataset. The aim of this procedure is to identify which reference population is most alike the extracted Eurasian ancestry.

### Visualization

PCA and outgroup F_3_ results were visualized in R version 3.6.1 using the libraries `ggplot2` [H, 2016]. Maps were created in R version 3.6.1 using `ggplot2` and the `sf, rnaturalearth` and `rnaturalearthdata` libraries. The kriging projection maps were generated in Surfer version 12.0.626 from Golden Software.

## Results

After quality control, removal of related individuals, and down-sampling to a maximum of 30 individuals per population the dataset consisted of 2066 individuals from 101 population groups and 199 422 SNPs. Note that some populations are represented multiple times from different original publications, resulting in a total of 97 unique populations.

### Population Structure in Northeast Africa

General population structure inferences were performed using PCA and ADMIXTURE on a dataset pruned of SNPs in LD (85 529 SNPs remaining). From both analyses, it is evident that the first division in the data set is between Africans and non-Africans, and it is clear that North- and East-Africans have a much larger proportion of shared ancestry with Eurasian groups than do other African groups (Supplementary Figure 6 and K=3 in Supplementary Figure 2). The first six principal components are shown in Supplementary Figure 5 to 19.

The output from Admixture shown in Figure 1 (for full analysis see Supplementary Figure 2) captures similar patterns to the PCA analysis, with the first separations being between major geographic regions, with North and East Africa showing closer association with Eurasian populations than groups from other parts of Africa does. The K with the lowest cross-validation error was K=13, see Supplementary Figure 3. The first division in the dataset, K=2 is between African and Non-African ancestry, whilst K=3 separates out East Asians (brown) and European ancestry (pink) – maximized in the TSI at 98.9%. The K=4 (purple) is maximized in the Ju’hoansi at 96.5% and seems to represent hunter-gather associated ancestry as it is also high in the RHGs. This purple component reaches 100% in the Ju’hoansiat K = 5 and above. It is at K=5 that East African groups break away from other African groups via affinity to the black component, which is maximized in the Nuba at 80.1%. Of particular interest for the present investigation is also the light green component that emerges is K=7 it is maximized in the Sabue population at 88%. It is also here that Sudanese populations separate from other East African groups that move from the Black to the light green (lime) component. The Sabue is one of the few remaining hunter-gatherer groups in East Africa today and they share genetic continuity with earlier hunter-gatherer ancestry from the region [Gopalan et al., 2022]. At K=11 another East African component appears maximized in the Somali populations and as such might represent Cushitic related ancestry.

As K=14 separates the Levantine populations from the Arabic populations, and distinguishing between these ancestries was of primary concern for the study we opted to visualize these two components further using a Kriging interpolation across the study area, Figure 1. These two ancestries were the dark blue component maximized in the Lebanese Druze and the blush pink component highest in the Qatari. To further investigate the historic patterns of gene flow, migrations, and which barriers to migrations are evident across the region of interest interest we used the FEEMS software package [Marcus et al., 2021], Figure 1 D.

In our Principal Component (PC) analyses, the first PC differentiates between African and non-African groups, with East Asians being furthest away from Africans, top left of Supplementary Figure 5. From this component, several African populations fall on the cline between African and non-African variation, in particular, this is North Africans, such as the Egyptians and populations from Chad who are known to have Eurasian admixture [Haber et al., 2016]. PC 3, in Supplementary Figure 6, explains the variation between Khoe-San groups at one extreme and West African groups on the other, with the RHGs on the same cline as Khoe-San groups. We also observe a grouping according to linguistics where Omotic, Afro-Asiatic, and Nilo-Saharan speakers separate from each other. East African groups position themselves here on the diagonal between PC 1 and PC 3, with the Ari, Sandawe, and Sabue populations forming their own cluster in the direction of the Khoe-San. In this projection, we can also see the shared genetic ancestry of East African, North African, and Middle Eastern groups with Eurasians. The Middle Eastern populations show a clear association with other Eurasian groups except for the East Asians. The plot of PC 1 vs PC 4, Supplementary Figure 7, highlights the Middle East’s position as a gateway between Africa and the rest of the world as Middle eastern individuals fall across the spectrum between the two extremes on PC 1. UMAP was also performed in the genotype information in our dataset, see Supplementary Figure 20. This analysis produces two larger clusters of populations, one consisting of West African groups, Eastern Bantu speakers, the Saharan speakers, the Nuer, Dinka and Shiluk from Sudan. The other cluster contains mainly Middle Eastern populations and Ethiopians, as well as the remaining Sudanese populations.

### Determination of Eurasian sources through local ancestry estimation

To identify distinct ancestries in East African populations, we employed MOSAIC [Salter-Townshend and Myers, 2019]. We wanted to identify patterns of local ancestries and determine which of our reference populations were the best proxies for the different genetic components. In particular, we were interested in the “non-African” or rather Eurasian segments of the genomes. Following the approach from previous studies, we wanted to isolate these Eurasian genetic segments and analyze them in isolation [Vicente et al., 2019], [Molinaro et al., 2019].

Thirty-five East African and Northeast African populations were chosen as target populations to analyze. For location and linguistic groups of these target populations see Supplementary Figure 4.

As has been shown in previous studies, and indicated by our demographic inference Figure 1 and Supplementary Figure 6, there are generally four main components of modern-day East African genetic ancestry [Prendergast et al., 2019, Wang et al., 2020]. Namely, basal East African hunter-gatherer ancestry, Sudanese/Nilotic ancestry, Eurasian ancestry, and West African ancestry associated with Bantu speakers. Since the aim of this study is to identify the best proxy for the source of the Eurasian ancestry of the Northeast African populations, we constructed a scenario where we could use these four ancestral sources to paint the haplotype of our chosen target populations using MOSAIC [Salter-Townshend and Myers, 2019]. We set up an initial scenario to try and identify the best Eurasian source to use for further analyses. In this scenario we used the following populations; To represent the basal East African hunter-gatherer ancestry we chose the Sabue as they are the modern-day population which best represents ancient East African hunter-gatherer ancestry [Scheinfeldt et al., 2019, Gopalan et al., 2022], see also Supplementary Figure 6 and 2. The Sabue have been referred to by many different name in the literature, for instance Shabo, Chabu, we have opted to use the name used in the original publication of our data [Scheinfeldt et al., 2019]. The CEU (Utah residents with Northern and Western European ancestry) population from the 1000 genomes consortium was chosen as a stand-in for general Eurasian ancestry. The Sudanese Dinka was chosen to represent the Sudanese ancestry since it is the group that defines the black component in the ADMIXTURE analysis associated with Sudanese ancestry. The Yoruba (YRI) was used as a proxy for West African Niger-Congo and Bantu-speaker-associated ancestry. We then used these four populations (CEU, YRI, Dinka, and Sabue) as sources in a 3-way admixture scenario in MOSAIC and extracted the called genotypes that were assigned to the CEU-like ancestry with a probability of 80% or more. The closest affinity of each constructed ancestry was determined by F_st_ tests against the four source populations. This resulted in regions of each East African individual’s genome that is more closely associated with a Eurasian ancestry than with the other ancestries. For information about the methodology see Materials and method on Local ancestry inference.

This non-African part of the genomes was then compared using outgroup F_3_ to Eurasian references populations with the Ju’hoansi as outgroup (target REF Ju’hoansi). A higher value of the outgroup F_3_ test indicates a smaller genetic distance between the in-groups compared to the outgroup. The San group Ju’hoansi was chosen based on the Admixture results, Figure 1, as it defines the dark purple component at 100 % for K’s *>* 5 and as previous studies had shown them to be the least admixed modern day Khoe-San group [Schlebusch et al., 2012]. The F_3_ outgroup test thus identified the population that is the most similar to the Eurasian fraction of the Northeast African target populations, see the top three in Table 1 and top five in Supplementary Table 2.

**Table 1:**
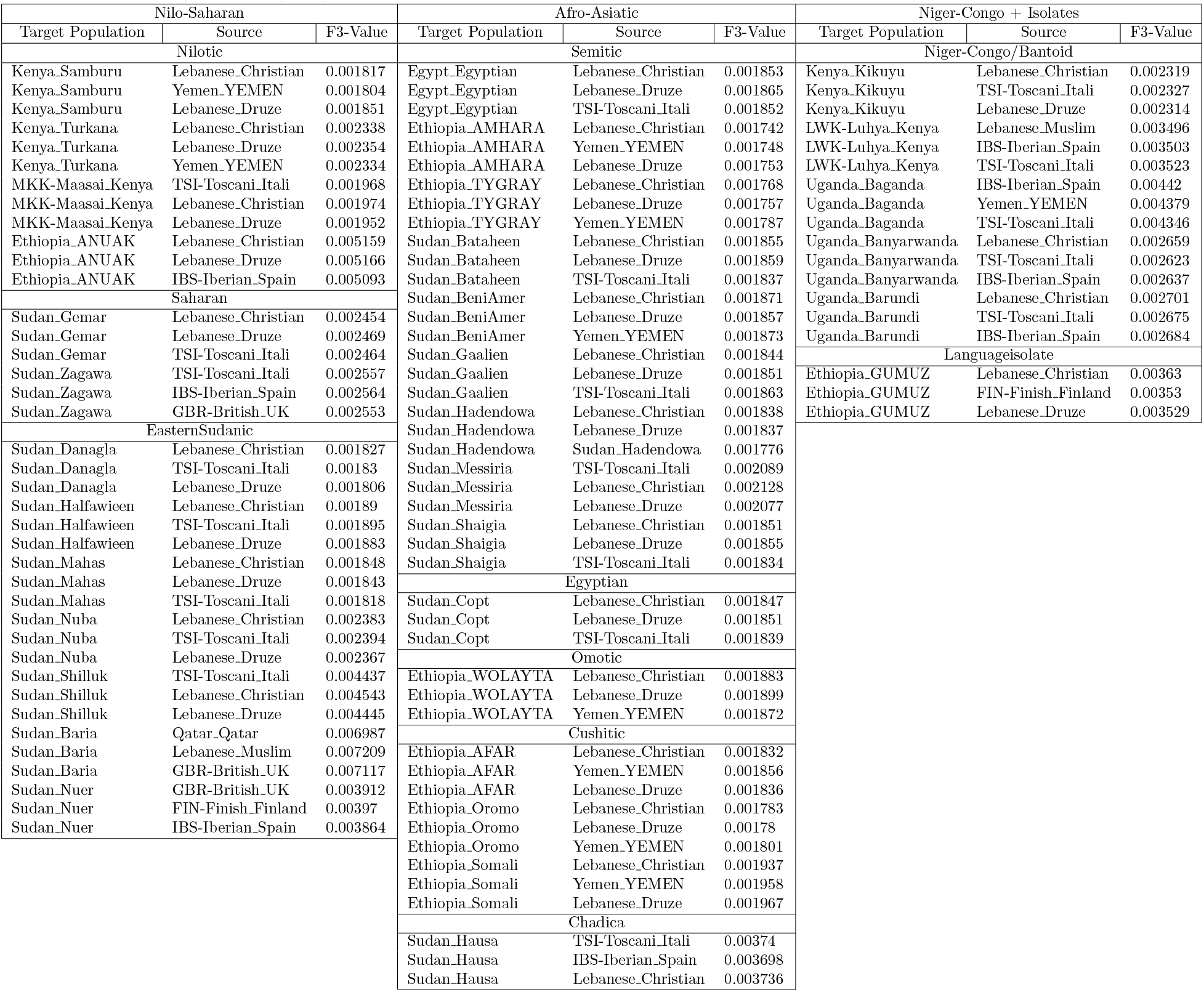
F_3_ outgroup result grouped by language family of the target populations, top 3 hits show. The F_3_ outgroup was calculated for the most Eurasian like ancestry for each target population in the following manner: Target Source Ju’hoansi

| Nilo-Saharan |  |  | Afro-Asiatic |  |  | Niger-Congo + Isolates |  |  |
| --- | --- | --- | --- | --- | --- | --- | --- | --- |
| Target Population | Source | F3-Value | Target Population | Source | F3-Value | Target Population | Source | F3-Value |
| Nilotic |  |  | Semitic |  |  | Niger-Congo/Bantoid |  |  |
| Kenya_Samburu | Lebanese_Christian | 0.001817 | Egypt_Egyptian | Lebanese_Christian | 0.001853 | Kenya_Kikuyu | Lebanese_Christian | 0.002319 |
| Kenya_Samburu | Yemen_YEMEN | 0.001804 | Egypt_Egyptian | Lebanese_Druze | 0.001865 | Kenya_Kikuyu | TSI-Tosceni_Itali | 0.002327 |
| Kenya_Samburu | Lebanese_Druze | 0.001851 | Egypt_Egyptian | TSI-Tosceni_Itali | 0.001852 | Kenya_Kikuyu | Lebanese_Druze | 0.002314 |
| Kenya_Turkana | Lebanese_Christian | 0.002338 | Ethiopia_AMHARA | Lebanese_Christian | 0.001742 | LWK-Luhya_Kenya | Lebanese_Muslim | 0.003496 |
| Kenya_Turkana | Lebanese_Druze | 0.002354 | Ethiopia_AMHARA | Yemen_YEMEN | 0.001748 | LWK-Luhya_Kenya | IBS-Iberian_Spain | 0.003503 |
| Kenya_Turkana | Yemen_YEMEN | 0.002334 | Ethiopia_AMHARA | Lebanese_Druze | 0.001753 | LWK-Luhya_Kenya | TSI-Tosceni_Itali | 0.003523 |
| MKK-Maasai_Kenya | TSI-Tosceni_Itali | 0.001968 | Ethiopia_TYGRAY | Lebanese_Christian | 0.001768 | Uganda_Baganda | IBS-Iberian_Spain | 0.00442 |
| MKK-Maasai_Kenya | Lebanese_Christian | 0.001974 | Ethiopia_TYGRAY | Lebanese_Druze | 0.001757 | Uganda_Baganda | Yemen_YEMEN | 0.004379 |
| MKK-Maasai_Kenya | Lebanese_Druze | 0.001952 | Ethiopia_TYGRAY | Yemen_YEMEN | 0.001787 | Uganda_Baganda | TSI-Tosceni_Itali | 0.004346 |
| Ethiopia_ANUAK | Lebanese_Christian | 0.005159 | Sudan_Bataheen | Lebanese_Christian | 0.001855 | Uganda_Banyarwanda | Lebanese_Christian | 0.002659 |
| Ethiopia_ANUAK | Lebanese_Druze | 0.005166 | Sudan_Bataheen | Lebanese_Druze | 0.001859 | Uganda_Banyarwanda | TSI-Tosceni_Itali | 0.002623 |
| Ethiopia_ANUAK | IBS-Iberian_Spain | 0.005093 | Sudan_Bataheen | TSI-Tosceni_Itali | 0.001837 | Uganda_Banyarwanda | IBS-Iberian_Spain | 0.002637 |
| Saharan |  |  | Sudan_BeniAmer | Lebanese_Christian | 0.001871 | Uganda_Barundi | Lebanese_Christian | 0.002701 |
| Sudan_Gemar | Lebanese_Christian | 0.002454 | Sudan_BeniAmer | Lebanese_Druze | 0.001857 | Uganda_Barundi | TSI-Tosceni_Itali | 0.002675 |
| Sudan_Gemar | Lebanese_Druze | 0.002469 | Sudan_BeniAmer | Yemen_YEMEN | 0.001873 | Uganda_Barundi | IBS-Iberian_Spain | 0.002684 |
| Sudan_Gemar | TSI-Tosceni_Itali | 0.002464 | Sudan_Gaalien | Lebanese_Christian | 0.001844 | Languageisolate |  |  |
| Sudan_Zagawa | TSI-Tosceni_Itali | 0.002557 | Sudan_Gaalien | Lebanese_Druze | 0.001851 | Ethiopia_GUMUZ | Lebanese_Christian | 0.00363 |
| Sudan_Zagawa | IBS-Iberian_Spain | 0.002564 | Sudan_Gaalien | TSI-Tosceni_Itali | 0.001863 | Ethiopia_GUMUZ | FIN-Finish_Finland | 0.00353 |
| Sudan_Zagawa | GBR-British_UK | 0.002553 | Sudan_Hadendowa | Lebanese_Christian | 0.001838 | Ethiopia_GUMUZ | Lebanese_Druze | 0.003529 |
| EasternSudanic |  |  | Sudan_Hadendowa | Lebanese_Druze | 0.001837 |  |  |  |
| Sudan_Danagla | Lebanese_Christian | 0.001827 | Sudan_Hadendowa | Sudan_Hadendowa | 0.001776 |  |  |  |
| Sudan_Danagla | TSI-Tosceni_Itali | 0.00183 | Sudan_Messiria | TSI-Tosceni_Itali | 0.002089 |  |  |  |
| Sudan_Danagla | Lebanese_Druze | 0.001806 | Sudan_Messiria | Lebanese_Christian | 0.002128 |  |  |  |
| Sudan_Halfawieen | Lebanese_Christian | 0.00189 | Sudan_Messiria | Lebanese_Druze | 0.002077 |  |  |  |
| Sudan_Halfawieen | TSI-Tosceni_Itali | 0.001895 | Sudan_Shaigia | Lebanese_Christian | 0.001851 |  |  |  |
| Sudan_Halfawieen | Lebanese_Druze | 0.001883 | Sudan_Shaigia | Lebanese_Druze | 0.001855 |  |  |  |
| Sudan_Mahas | Lebanese_Christian | 0.001848 | Sudan_Shaigia | TSI-Tosceni_Itali | 0.001834 |  |  |  |
| Sudan_Mahas | Lebanese_Druze | 0.001843 | Egyptian |  |  |  |  |  |
| Sudan_Mahas | TSI-Tosceni_Itali | 0.001818 | Sudan_Copt | Lebanese_Christian | 0.001847 |  |  |  |
| Sudan_Nuba | Lebanese_Christian | 0.002383 | Sudan_Copt | Lebanese_Druze | 0.001851 |  |  |  |
| Sudan_Nuba | TSI-Tosceni_Itali | 0.002394 | Sudan_Copt | TSI-Tosceni_Itali | 0.001839 |  |  |  |
| Sudan_Nuba | Lebanese_Druze | 0.002367 | Omotic |  |  |  |  |  |
| Sudan_Shilluk | TSI-Tosceni_Itali | 0.004437 | Ethiopia_WOLAYTA | Lebanese_Christian | 0.001883 |  |  |  |
| Sudan_Shilluk | Lebanese_Christian | 0.004543 | Ethiopia_WOLAYTA | Lebanese_Druze | 0.001899 |  |  |  |
| Sudan_Shilluk | Lebanese_Druze | 0.004445 | Ethiopia_WOLAYTA | Yemen_YEMEN | 0.001872 |  |  |  |
| Sudan_Baria | Qatar_Qatar | 0.006987 | Cushitic |  |  |  |  |  |
| Sudan_Baria | Lebanese_Muslim | 0.007209 | Ethiopia_AFAR | Lebanese_Christian | 0.001832 |  |  |  |
| Sudan_Baria | GBR-British_UK | 0.007117 | Ethiopia_AFAR | Yemen_YEMEN | 0.001856 |  |  |  |
| Sudan_Nuer | GBR-British_UK | 0.003912 | Ethiopia_AFAR | Lebanese_Druze | 0.001836 |  |  |  |
| Sudan_Nuer | FIN-Finish_Finland | 0.00397 | Ethiopia_Oromo | Lebanese_Christian | 0.001783 |  |  |  |
| Sudan_Nuer | IBS-Iberian_Spain | 0.003864 | Ethiopia_Oromo | Lebanese_Druze | 0.00178 |  |  |  |
|  |  |  | Ethiopia_Oromo | Yemen_YEMEN | 0.001801 |  |  |  |
|  |  |  | Ethiopia_Somali | Lebanese_Christian | 0.001937 |  |  |  |
|  |  |  | Ethiopia_Somali | Yemen_YEMEN | 0.001958 |  |  |  |
|  |  |  | Ethiopia_Somali | Lebanese_Druze | 0.001967 |  |  |  |
|  |  |  | Chadica |  |  |  |  |  |
|  |  |  | Sudan_Hausa | TSI-Tosceni_Itali | 0.00374 |  |  |  |
|  |  |  | Sudan_Hausa | IBS-Iberian_Spain | 0.003698 |  |  |  |
|  |  |  | Sudan_Hausa | Lebanese_Christian | 0.003736 |  |  |  |

The outgroup F_3_ analysis provided us with the best Eurasian source population to use for each of our Northeast African target populations. We then re-ran the MOSAIC analysis above but instead using this best source instead of the CEU. We refer to this dataset and approach as the “best by F_3_”.

In addition to the above scenario, using CEU as a proxy and then identifying the best ancestral source through F-statistics, we also investigated some more unconstrained scenarios. In these scenarios, we kept YRI, Sabue, and Dinka as the three African populations and then varied the Eurasian sources. We tried both providing one Eurasian population at a time as well as providing two Eurasian populations at the same time. These scenarios were repeated under a 2-way, 3-way, and 4-way admixture scenario, that is using two, three and four ancestral sources with the four or five reference populations respectively. All 35 target populations were investigated under these differing permutations. Since evaluating the best model can be nontrivial and require lots of manual curation we opted to use MOSAIC’s R^2^ metric (genomic fit) to evaluate the best model. In general, the simpler models performed better, all of the 2-way scenarios outperformed their equivalent (using the same populations) 3-way and 4-way admixture scenarios. Though using two Eurasian populations as sources outperformed a single source. These R^2^ values can be found in Supplementary Table 1. Thus these scenarios resulted in a dataset with a 2-way admixture scenario with YRI, Sabue, Dinka, and in addition two Eurasian populations as the sources for these ancestries. This dataset is referred to as the “Best by R^2^” or simply “R^2^” dataset in the rest of the study. These runs thus produces two inferred ancestral sources, one Non-African and one African. Five of the target populations did not generate a Non-African source as the closest fit determined by Fst, these were LWK-Luhya Kenya, Sudan Baria, Sudan Hausa, Sudan Nuer, and Uganda Baganda thus none of these populations are shown in the best by R^2^ analysis. Ancestry tract length distribution plots for both of these datasets was generated and are available on request.

### Dating Eurasian admixture dating in Northeast Africa

For both the F_3_ outgroup based approach and the R^2^ approach above, we determined the admixture date (in generations) from MOSAIC’s co-ancestry curves for the Eurasian-like constructed ancestry. The results of this dating can be seen in Figure 2 with the best source based on F_3_ in A, and D and the dates based on the runs with the highest R^2^ value in B and E. Given a generation time of 29 years this gives a time span of 72.5 years ago for the Nilotic speaking Anuak to 1027 years ago for the Cushitic speaking Afar, both from Ethiopia for the best by R^2^ dataset [Fenner, 2005]. In the best by F_3_ dataset the range is smaller with 84 years for the Eastern Sudanic speaking Nuer (South Sudan) to 940 for the Semitic speaking Bataheen (Sudan).

**Figure 2:**
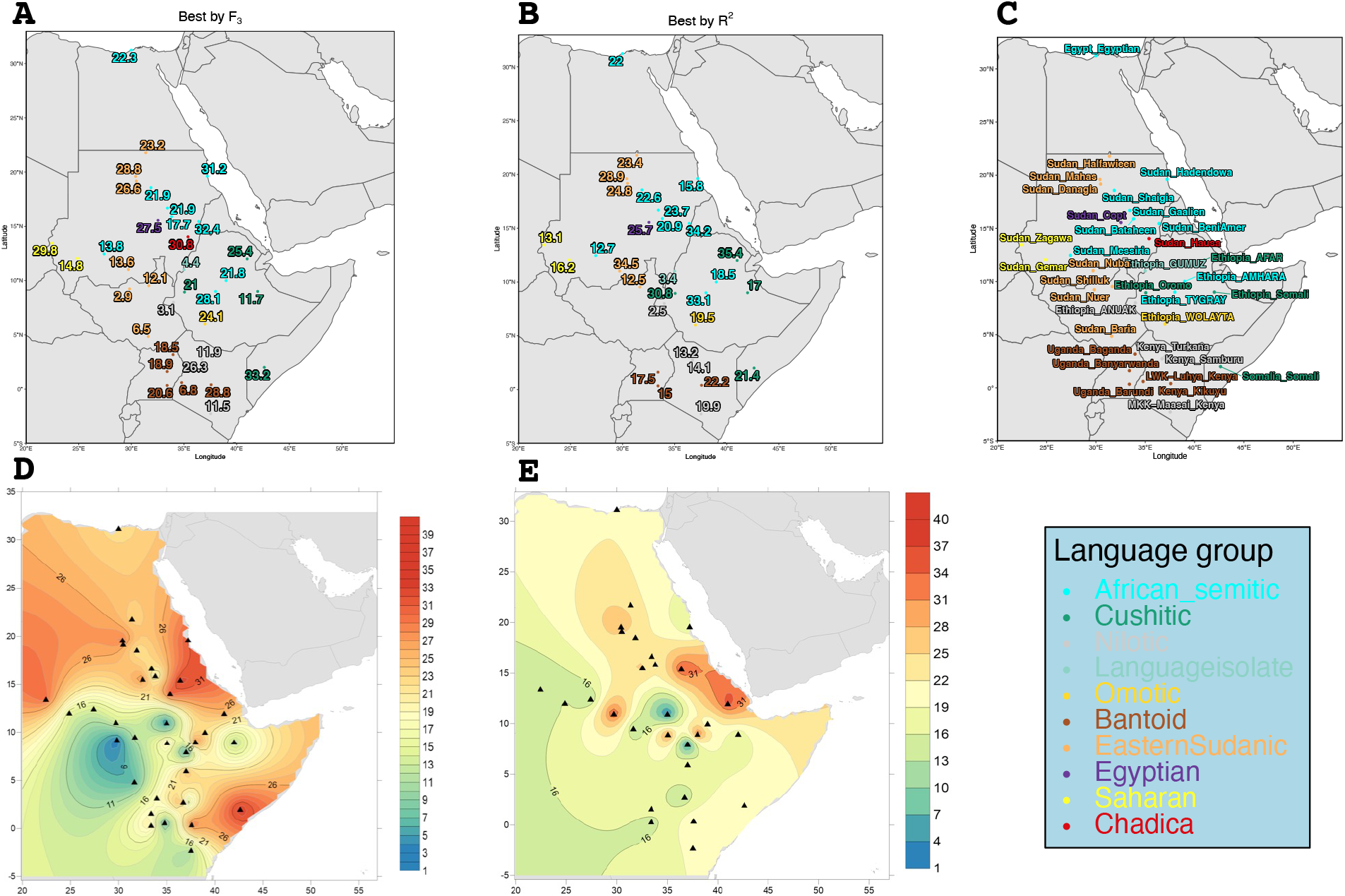
Admixture dating in generation for the most Eurasian like ancestry from MOSAIC. A and D contain data for the best source as determined by F_3_ whilst B and E illustrate the dataset determined by the best on R^2^ value. A and B is the admixture date in generations, C is the target populations locations, D and E is the same data but plotted over the study area surface using Kriging interpolation. The numbers here represent the major breaks (black lines). Note that some populations did not find a Eurasian source in the best by R^2^ runs and thus do not have date.

To visualize the correlation between linguistic classification and the inferred admixture date we generated dot plots of the dates per linguistic group as well as the larger linguistic families, see Figure 3.

**Figure 3:**
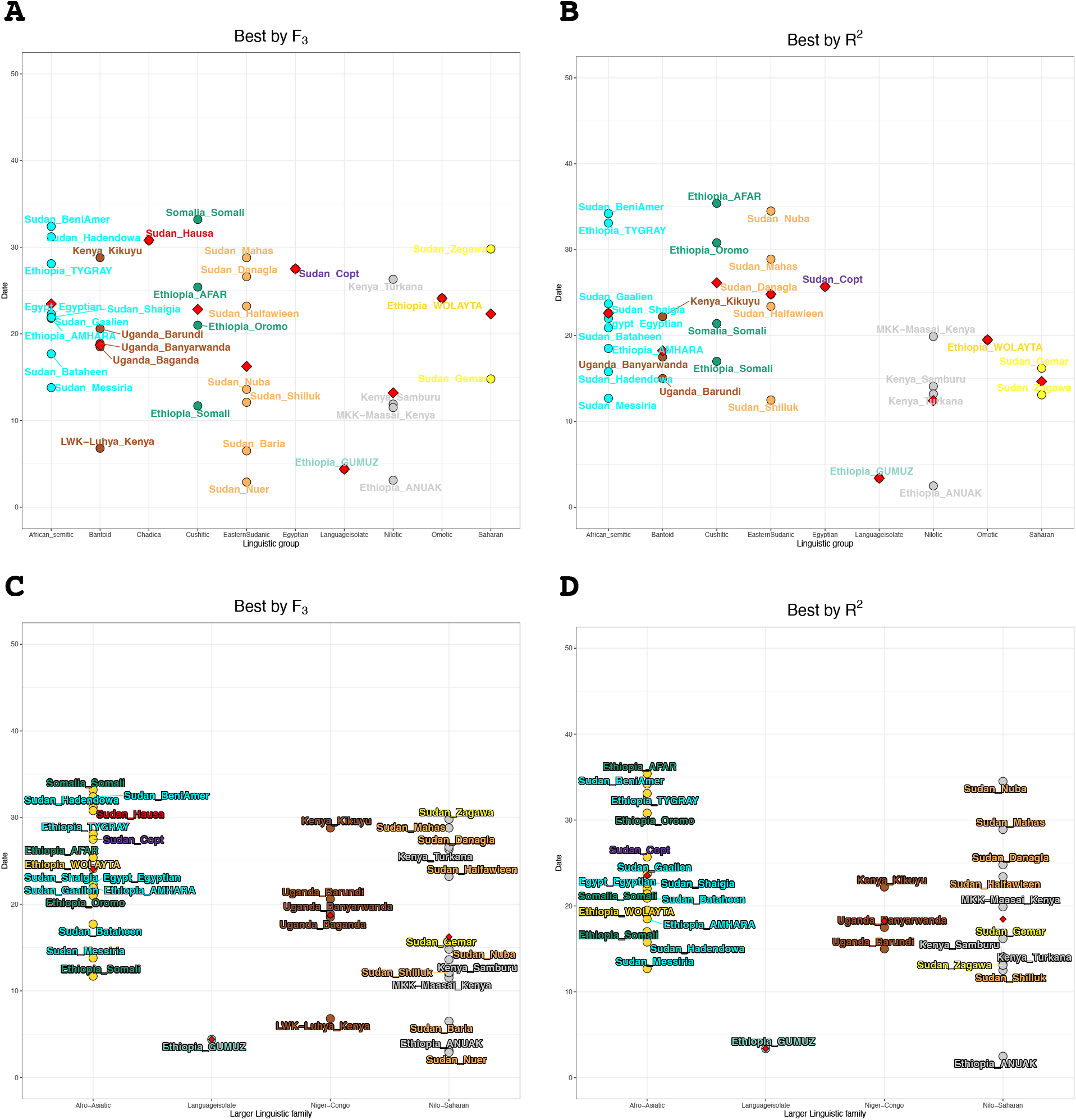
Dot plot representations of the admixture dating in generation for the most Eurasian-like ancestry from MOSAIC. A and C contain data for the best source as determined by F_3_ whilst B and D illustrate the dataset determined by the best on R^2^ value. A and B are per smaller linguistic classification whilst C and D show the same data but divided into linguistic family. The red triangle represents the mean value.

We compared these dates to the categorical information we had about the populations, that is Country, Linguistic group (e.g. Semitic), or larger Linguistic family (e.g. Afro-Asiatic) using a two-way ANOVA, Supplementary Table 4. Only larger linguistic families correlated significantly with the detected admixture dates for the best by F_3_ dataset, Supplementary Table 4 A. The same pattern where the lowest p-value is observed for the larger linguistic family is true also for the R^2^ dataset but without reaching significance, Supplementary Table 4 B. We also test whether there was a spatial correlation to the admixture dates. This was done by comparing the great circle distance between Tel Aviv (a coastal location in the Levant) as well as Sanaa (the capital of Yemen) and each population’s sampling location, Supplementary Figure 21. For the R^2^ values by each linguistic family see Supplementary Table 3. For the distance from Tel Aviv we find a low but significant correlation for both datasets, R_2_ of 0.088 for the best by F_3_ dataset and 0.067 for the best by R^2^ data. We find weaker, but significant, support for the distance from Sanaa in both datasets R^2^ 0.048 (p = 0.001) for the best by F_3_ dataset and R_2_ 0.031 (p = 0.002), Supplementary Figure 21.

As there was some discrepancies between the two dating approaches we decided to compare the dates to each other. To do that we plotted the dates received from the F_3_ dataset against the R^2^ dataset, see Figure 4. Then we performed linear regression on the data. This resulted in a correlation (R^2^) of 0.3093 with a p-value of 0.001417. The majority of target populations falls within the 95% confidence interval, gray area in Figure 4.

**Figure 4:**
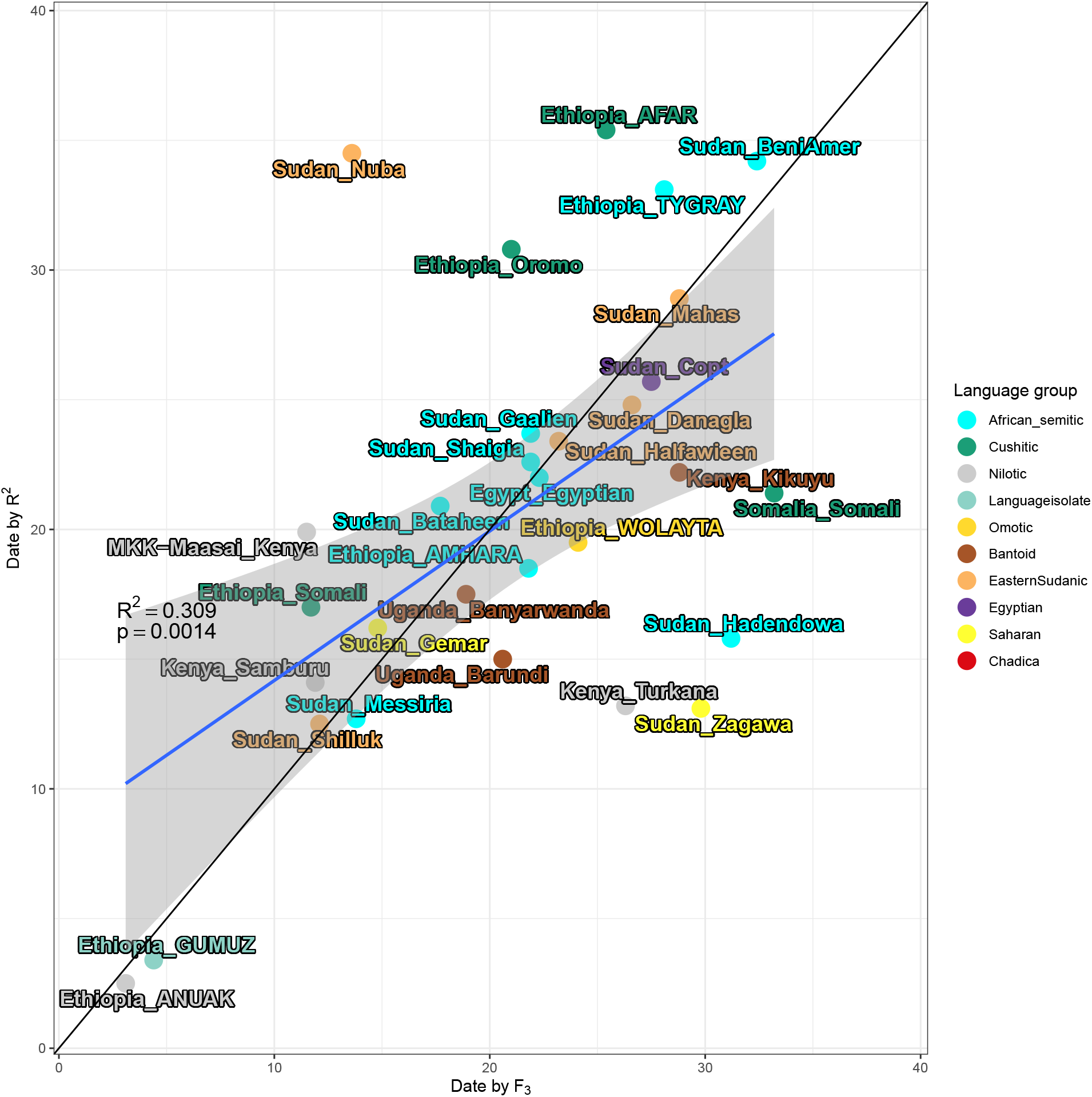
Linear regression (blue line) comparing the admixture dates of the most Eurasian-like constructed ancestry from the best by F_3_ to the best by R^2^ dataset. The greyed area represents the 95% confidence interval. Black line is X=Y, i.e. same date in both approaches.

## Discussion

In this study, we investigate the patterns of different genetic ancestries in Northeast African populations, with a focus on Eurasian back migrations. We inferred population structure using both global and local ancestry methods. Using the local ancestry method MOSAIC we identify regions of Northeast African populations genomes with Eurasian ancestry. We also attempt to date this admixture. Northeast Africa is a region of complex history spanning back thousands of years. The expansions and contractions, rise and fall of states, kingdoms and empires across the region have had a major impact on the formation, dissolution, and current distributions of the sampled communities in this study. We therefore recognize that the groups included in this study are modern-day populations that were created by introgression/interaction/assimilation events in the past and should not be seen unchanged entities that represent exact past distributions of groups. Rather we start from the modern-day groups and try to infer some patterns of past interactions through analysing their genomes.

Population structure inference illustrates the complex genetic history of Northeast African populations. Larger patterns of genetic associations between many of the world’s distinct human lineages are reflected in Northeast African genomes. The hunter-gatherer’s ancestry highlights the deep history of the region and its people and that this ancestry remains within the East African populations. The southern part of the region has a closer genetic affinity to West African groups, a result of the Bantu expansion and several of these populations also speak Bantu languages today. That the Bantu expansion did not continue further into the region could be a result of geographical barriers such as the Ethiopian Highlands and the dry regions of the Horn of Africa, indicated by our FEEMS analysis 1 D or as suggested by Hollfelder et al. [2017] that the Northeast African Nilotic speaking herders (such as the Dinka and Nuer), who have remained relatively isolated from other groups, could have formed a buffer against the Bantu expansion continuing further into Northeast Africa. The interpretation of the dating for Nilotic speakers from East Africa, the Maasai, Turkana, and Samburu is itself complex since they only relatively recently reached their current day distributions through expansions from Sudan and Uganda within the last few centuries [Spear and Waller, 1993]. Our dating thus indicate that they picked up their Eurasian ancestry before this migration.

Eurasian admixture have had a large influence on the genomes of Northeast African groups. The Egyptians and Sudanese Copt populations for instance are genetically very similar to Middle Eastern groups rather than to other African populations. The pattern is true also for the rest of North Africa, though not investigated here. This pattern is ancient, being present as early as at least 15 000 years ago [van de Loosdrecht et al., 2018]. Further south in the region we continue to see the impact of past Eurasian admixture. Northeast African populations’ positions in the PCA plots are being drawn towards Eurasian populations on PC 1, Supplementary Figure 6. Admixture analyses recapitulate this pattern where Northeast African groups share the blue component maximized in Middle Eastern groups at K=7, Figure 2.

The Copts look genetically similar to the Egyptians from Cairo, see Figure 1 A and C, this is not unsurprising given that the Copts arrived in Sudan around 200 years ago from Egypt and seems to have lived relatively isolated since then [Hollfelder et al., 2017]. Our admixture date for the Copts was inferred to be 27.5 for the F_3_ analysis and 25.7 for the R^2^ and around 22 generations for the Egyptians. Thus this admixture took place around the 14:th century.

Previous studies that investigated this Northeast Africa, Ethiopia in particular, observed clustering based on linguistic families [Pagani et al., 2012, Hellenthal et al., 2021, Hollfelder et al., 2017]. This pattern is recapitulated in our analysis, both from the population inference methods as well as the admixture dating. Looking at PC 1 vs PC 3 in Supplementary Figure 6, the Omotic speaking Ari populations form their own small cluster, a reflection of their segregated position within Ethiopian society [Pagani et al., 2012].

Another part of the complex East African history is the relationships between ancient hunter-gather groups. In PC 3 there is a cline between the Khoe-San groups and RHGs at one end to East Africans at the other, this is most clearly seen in PC 3 vs PC 5 in Supplementary Figure 15. This cline is similar to what was found in studies using aDNA [Skoglund et al., 2017, Prendergast et al., 2019] and could be a reflection of East African hunter-gather ancestry. Similarly in the Admixture analysis, we see a small proportion of the maroon proportion that is maximized in Khoe-San groups and RHGs at K=4, Supplementary Figure 2. The ancient hunter-gather ancestry in modern-day East African groups, Sudanese excluded, is consistent with the analyses based on the Mota genome [Gallego Llorente et al., 2015].

The Gumuz and Sabue have both been suggested to retain a high degree of ancient East African hunter gatherer ancestry, [Pagani et al., 2012, Scheinfeldt et al., 2019, Gopalan et al., 2022] and our demographic analyses indicates a high degree of similarity between the two populations, such as the ADMIXTURE results where the Gumuz and Sabue have a high proportion of the dark green component. This dark green component is retained even when the other East African groups switch over to the red component at K=13, Supplementary Figure 2, previous studies also observed this pattern for both the Sabue and the Gumuz [Gopalan et al., 2022]. The Cushitic speakers remain quite similar throughout the different K’s, with the major proportions being the Middle Eastern maximized light blue and the East African associated dark green being the two main components. The Gumuz and Anuak display very little Eurasian admixture, and given that the date that we infer is only a few generations ago, it could be that they have received this Eurasian admixture through secondary admixture with another neighbouring group a few generations ago, or that it’s an effect of recent or ongoing admixture.

Dating the admixture of different groups with each other is of great interest to population geneticists. Having a date for when the mixing of two groups occurred allows us to incorporate other types of independent evidence into our analyses, such as written or oral history or linguistic information, thus a big part of the effort in this study and discussion is spent on our inferred dates of admixture. The “best by F_3_” analysis is our attempt to propose a scenario that best fits the previously known genetic history of the region, whilst the “best by R^2^” analysis, were we picked a 3-way admixture event and groups based on the genomic fit (R^2^), is intended as a less constrained scenario for picking out Eurasian ancestry in Northeast African groups. The Sudanese data in our study is mainly from Hollfelder et al. [2017] who also investigated the time and sources of admixture in Sudanese populations. Our findings are generally in agreement, particularly for the Eurasian admixture dates that is the primary focus of our study. Hollfelder et al. [2017] investigated a simpler admixture scenario with only two putative sources, namely the Sudanese Nuer and the TSI (Tuscan) to represent the admixture of a Sudanese basal population with a Eurasian source. This is most similar to our R^2^ approach in which we picked the scenario with the best genomic fit (R^2^) and for two Eurasian sources in each run and then picked the two Eurasian sources that produced the best genomic fit (R^2^ value). As shown in Figure 4 the two approaches results in similar dates for most of the populations. There is a linear relationship between the inferred date in the 3-way best by F_3_ scenario and the best by R^2^ scenario, of 0.3093 and p-value = 0.001417. Most of the populations that fall outside the 95% SE confidence interval do fall on the X=Y line instead. Notable exceptions are the Sudanese Nuba and the Ethiopian Afar populations that have much older dates in the R^2^ scenario, and the Sudanese Zagawa and Hadendowa, who display the opposite pattern with much older date in the best by F_3_ scenario. Most other populations that deviate from either line do so with only a few generations. From our analyses there is nothing in particular that makes these four populations stand out from nearby populations such as in the ADMIXTURE or PCA. The MOSAIC metric such as R^2^, R_st_ etc are also in line with comparable populations. The Sudanese Nuba are known to be a relatively heterogeneous group [Spaulding, 2014], but that is not reflected in our analyses of population structure, see Figure 1 and Supplementary Figure 6.

The analysis using FEEMS recapitulates expected natural barriers to migration such as the Red Sea, the Gulf of Aden, the Persian Gulf, and the Sahara Desert. In addition to clear geographical barriers, we also see evidence of linguistic and cultural barriers. One obvious example is the low migration rate between the Ethiopian Somali and the other Ethiopian populations, and as expected high migration rate inferred between the different Somali groups. The Great Rift Valley forms a natural barrier across Ethiopia with highlands of on both sides of the rift. A previous study looking at Ethiopian genetics found significant association of genetic similarity to elevation, ethnicity and first language, and interestingly not second language nor religion [Ĺopez et al., 2021]. Along the Red Sea coast of Eritrea and Sudan, we find a region of high gene flow extending into northern Ethiopia and into the Great Rift Valley, Figure 1 D. This region corresponds well to the pink component in Figure 1 A and B which seems to represent Yemeni ancestry. It is also a region in which we infer some of the oldest inferred admixture dates. These observations, as well as the shared linguistics of South Semitic as South Semitic languages that are found in Yemen, Oman, Eritrea, and Ethiopia [Faber, 1980], indicate a close connection between Eritrea, and Ethiopia to the South of the Arabian peninsula and present-day Yemen. The Kingdom of Aksum (or the Aksumite Empire) Eastern Sudan, Northen Ethiopia, Eritrea and Djibouti to across the Red Sea into Yemen thrived between the 1:th and 7:th century AD, as trade along the Red Sea increased and the trade along the Nile decreased. Both Rome and Byzantium traded with the Indian Subcontinent and artefacts from these Kingdoms can be found at Aksumite sites, Finneran [2000], Sharp [1999]. The Semitic-speaking Ethiopian populations also group together with the Middle Eastern populations in the UMAP analysis, Supplementary Figure 20. These admixture event(s) could come as the result of the Red Sea trade. Aksum collapsed in the 8:th century as Islam started to expand and control over the Red Sea trade shifted to the Near East [Mitchell and Lane, 2013]. There is a clear divide between the Eastern Sudanic Semitic speaking groups and the South Sudanese groups as well as the Saharan-speaking Sudanese groups, both with regards to global ancestry as well as their inferred admixture dates for their Eurasian ancestries, which distinguishes them from the populations along the Red Sea coast.

Previous recent ancestry deconvolution studies pointed at Levantine sources for the Eurasian admixture in Northeast Africans rather than Arabic groups [Pagani et al., 2012, Molinaro et al., 2019]. We find that the pattern to be more complex than that with differing source populations in differing regions, see Figure 1 B and C as well as Table 1. The Lebanese Christians and the Yemeni populations are both among the top pick of sources for many populations in our 1 outgroup analysis. Admixture analyses suggest Arabian source populations rather than Levantine contributions, Figure 1 B and C. Dongola had been the capital of the Nubian Kingdom and the fall of Dongola in 317 to Mameluke forces meant the start of Arab and Islamic dominance south of the borders of Egypt. Many of the Semitic speakers in our dataset have their Eurasian admixture dated to this time – around 20+ generations ago. The exception is mainly the Southern Semitic speakers such as the Beni-Amer and Tygray whose dates are slightly older at around 30 generations ago. Around 30 generations ago is also the inferred dates for the Ethiopian Cuschitic speaking Afar and Oromo (though Oromo had a generation time of 21 for the best by F_3_).

Major linguistic family was the only factor that was significant (and only for the best by F_3_) in our ANOVA test of the available categories, Supplementary Table 4. The linear regression analysis of distance from the Levant, Supplementary Figure 21 A and B, also produced a significant fit with a negative coefficient indicating more recent admixture dates further from the Levant – this is likely driven by the younger dates for the populations in and around South Sudan. The same pattern was observed when comparing the distance to Sanaa, albeit with a smaller slope of the line and larger p-values (Supplementary Figure 21 C and D).

One possible explanation for this phenomenon could be that populations with little or no previous Eurasian admixture would have their inferred admixture date effected more by recent Eurasian admixture than population that experienced larger admixture in the past. In other words, most, if not all, of the populations in this study have or have had admixture with populations from the Middle East during the Arab expansion, and this newer admixture is obscuring older admixture patterns. The groups with younger inferred dates in our analysis thus have less the of older admixture. This pattern is also evident in the ADMIXTURE analysis, as the populations around South Sudan are represented by the black component at K=5 and onward rather than the pink component that we find in most other North-East African groups, indicating their isolation and genetic homogeneity compared to other populations. Older events that also contributed to admixture patterns in Northeast Africa are those that we have alluded to previously such as linguistic stratification, the trade across the Red Sea and the fall of Aksum.

The fact that both approaches for admixture dating produced populations from the same country (or previous countries in the case of Sudan and South Sudan) that had the the most extreme difference in Eurasian admixture dating, highlights the heterogeneous nature of North-East African genetics and how little explanatory power country borders have on population structure. It is, not unexpectedly so, rather geographic, linguistic and cultural borders that explain the degree of genetic interconnectedness between groups.

### Final remarks

We found that identifying the impact of ancient events on populations was not feasible when the original pattern has been distorted or masked by subsequent admixture events. Our study thus points to that the distribution of Eurasian-like ancestry in Eastern and North-Eastern African populations is mostly an effect of more recent migrations (many of them recorded in historical texts) rather than ancient events related to the advent of pastoralism in the region at large. To fully explore the question of Eurasian admixture into Africa over larger timescales likely requires population-level aDNA, especially of the early East African hunter-gatherers such as Mota, and the various in-moving groups, including those containing Eurasain admixture. North-Eastern Africa is a vast region with complex histories of migrations and admixture. It was not possible to identify one source or origin of Eurasian admixture in the region, rather different populations have experienced admixture at different times, at varying degrees, and from different external sources. Although slight trends were observed linked to language grouping and geography, the overall pattern proved to be complex and specific to certain population groups. Previous studies have highlighted these events in distinct regions or countries in Northern and Eastern Africa, whilst we in this study have tried to combine them with a specific emphasis on the Eurasian admixture in modern-day populations.

## Acknowledgements

The computation and data handling were enabled by resources provided by the Swedish National Infrastructure for Computing (SNIC) at Uppmax partially funded by the Swedish Research Council through grant agreement no. 2018-05973. Authorized NIH Data Access Committee (DAC) granted data access to Carina Schlebusch for the controlled-access genetic data analysed in this study that were previously deposited by Scheinfeldt et al. 2019 in the NIH dbGAP repository (accession code phs001780.v1.p1; date of approval: 2019-05-17). For the genome-wide genotype data from the Patin et al. 2017 study (EGA accessory number EGAD00010001209), data access was granted via European GenomePhenome Archive (EGA) by the GEH Data Access Committee EGAC00001000139. This project was supported by funding to CS from the European Research Council (ERC) under the European Union’s Horizon 2020 research and innovation programme (grant agreement No. 759933). A special thanks to the author of MOSAIC Michael Salter-Townshend for discussion on how the best perform the ancestry deconvolution, to Carolina Bernhardsson for help with plotting and Cesar Fortes-Lima for help with the study design.

## Supplementary information and Figures

**Supplementary Table 1:** Top two Eurasian source populations identified by their haplotype fit to the genomes (R^2^ value from MOSAIC) for each target population.

| Target | Sources | R2 |
| --- | --- | --- |
| Ethiopia_AFAR | GIH-Gujarati_India & Lebanese_Christian | 0.7097513 |
| Ethiopia_AMHARA | Dubai_Dubai & GIH-Gujarati_India | 0.7170762 |
| Ethiopia_TYGRAY | GIH-Gujarati_India & Lebanese_Muslim | 0.7086051 |
| Ethiopia_WOLAYTA | FIN-Finish_Finland & GIH-Gujarati_India | 0.7322527 |
| Ethiopia_Oromo | FIN-Finish_Finland & Lebanese_Christian | 0.7271779 |
| Egypt_Egyptian | IBS-Iberian_Spain & Lebanese_Druze | 0.8125434 |
| Ethiopia_GUMUZ | Dubai_Dubai & Oman_Oman | 0.765788 |
| Ethiopia_ANUAK | Oman_Oman & SaudiArabia_SaudiArabia | 0.7631948 |
| Ethiopia_Somali | Dubai_Dubai & Iran_Iran | 0.6993737 |
| Somalia_Somali | GIH-Gujarati_India & Yemen_YEMEN | 0.7882802 |
| Kenya_Samburu | Dubai_Dubai & Oman_Oman | 0.7269168 |
| Kenya_Turkana | Dubai_Dubai & Yemen_YEMEN | 0.8463287 |
| Kenya_Kikuyu | Lebanese_Muslim & Yemen_YEMEN | 0.7181286 |
| MKK-Maasai_Kenya | SaudiArabia_SaudiArabia & TSI-Toscani_Itali | 0.7204982 |
| LWK-Luhya_Kenya | Oman_Oman & Qatar_Qatar | 0.75834 |
| Uganda_Baganda | Iran_Iran & Oman_Oman | 0.7625496 |
| Uganda_Banyarwanda | Lebanese_Muslim & Yemen_YEMEN | 0.7379781 |
| Uganda_Barundi | Lebanese_Muslim & TSI-Toscani_Itali | 0.787222 |
| Sudan_Baria | Dubai_Dubai & GIH-Gujarati_India | 0.7562114 |
| Sudan_Bataheen | GIH-Gujarati_India & Lebanese_Christian | 0.8134197 |
| Sudan_BeniAmer | GIH-Gujarati_India & Lebanese_Muslim | 0.6988863 |
| Sudan_Copt | IBS-Iberian_Spain & Lebanese_Christian | 0.697059 |
| Sudan_Danagla | IBS-Iberian_Spain & TSI-Toscani_Itali | 0.7615013 |
| Sudan_Gaalien | GBR-British_UK & Lebanese_Christian | 0.7974334 |
| Sudan_Gemar | Lebanese_Christian & Lebanese_Muslim | 0.8498585 |
| Sudan_Hadendowa | GIH-Gujarati_India & Iran_Iran | 0.8009134 |
| Sudan_Halfawieen | GIH-Gujarati_India & Lebanese_Christian | 0.7735757 |
| Sudan_Hausa | Dubai_Dubai & Oman_Oman | 0.7649491 |
| Sudan_Mahas | IBS-Iberian_Spain & TSI-Toscani_Itali | 0.7600216 |
| Sudan_Messiria | Iran_Iran & Lebanese_Druze | 0.8687931 |
| Sudan_Nuba | FIN-Finish_Finland & Lebanese_Druze | 0.8584353 |
| Sudan_Nuer | Oman_Oman & Yemen_YEMEN | 0.7484863 |
| Sudan_Shaigia | GIH-Gujarati_India & Yemen_YEMEN | 0.8093127 |
| Sudan_Shilluk | GIH-Gujarati_India & TSI-Toscani_Itali | 0.758663 |
| Sudan_Zagawa | Dubai_Dubai & Oman_Oman | 0.7174841 |

**Supplementary Table 2:**
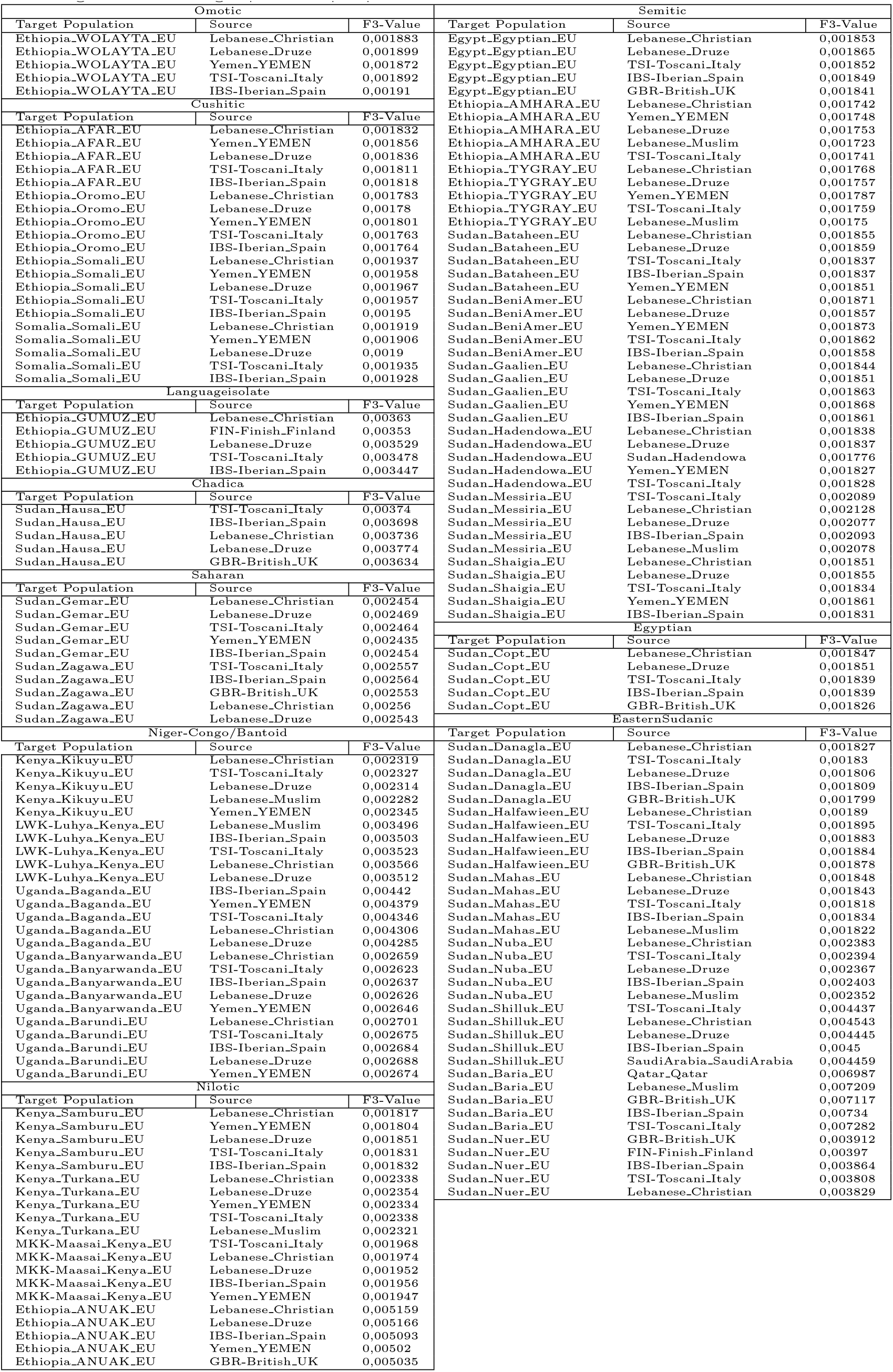
F_3_ outgroup result grouped by language family of the target populations, top 5 hits show. The F_3_ outgroup was calculated for the most Eurasian like ancestry for each target population in the following manner: Target Source Ju’hoansi.

**Supplementary Table 3:** R^2^ values of linear regression between admixture date and distance from Tel Aviv in the top row by each linguistic group. The Best by F_3_ dataset is in the leftmost columns while the Best by R^2^ dataset is on the right. Some linguistic groups have only one target population so no value whilst Saharan two populations which yields a R^2^ of 1.

| Distance from Tel Aviv |  |  |  |
| --- | --- | --- | --- |
| Best by $F_3$ | | Best by $F_2$ | |
| Linguistic group | $R^2$ | Linguistic group | $R^2$ |
| African_semitic | 0.002937986 | African_semitic | 0.010044898 |
| Cushitic | 0.303608661 | Cushitic | 0.383364613 |
| Nilotic | 0.138060608 | Nilotic | 0.967341737 |
| Languageisolate | - | Languageisolate | - |
| Omotic | - | Omotic | - |
| Bantoid | 0.003136976 | Bantoid | 0.006483384 |
| EasternSudanic | 0.830858222 | Egyptian | - |
| Egyptian | - | EasternSudanic | 0.033576907 |
| Saharan | 1.000000000 | Saharan | 1.000000000 |
| Chadica | - | Chadica | - |
| Distance from Sanaa |  |  |  |
| Best by $F_3$ | | Best by $R^2$ | |
| Linguistic group | $R^2$ | Linguistic group | $R^2$ |
| African_semitic | 0.02863027 | African_semitic | 0.04740316 |
| Cushitic | 0.26499817 | Cushitic | 0.43091650 |
| Nilotic | 0.16364704 | Nilotic | 0.98158939 |
| Languageisolate | - | Languageisolate | 0.00000000 |
| Omotic | - | Omotic | 0.00000000 |
| Bantoid | 0.06104717 | Bantoid | 0.42390308 |
| EasternSudanic | 0.82342297 | Egyptian | 0.00000000 |
| Egyptian | - | EasternSudanic | 0.07372633 |
| Saharan | 1.00000000 | Saharan | 1.00000000 |
| Chadica | - | Chadica | - |

**Supplementary Figure 1:**
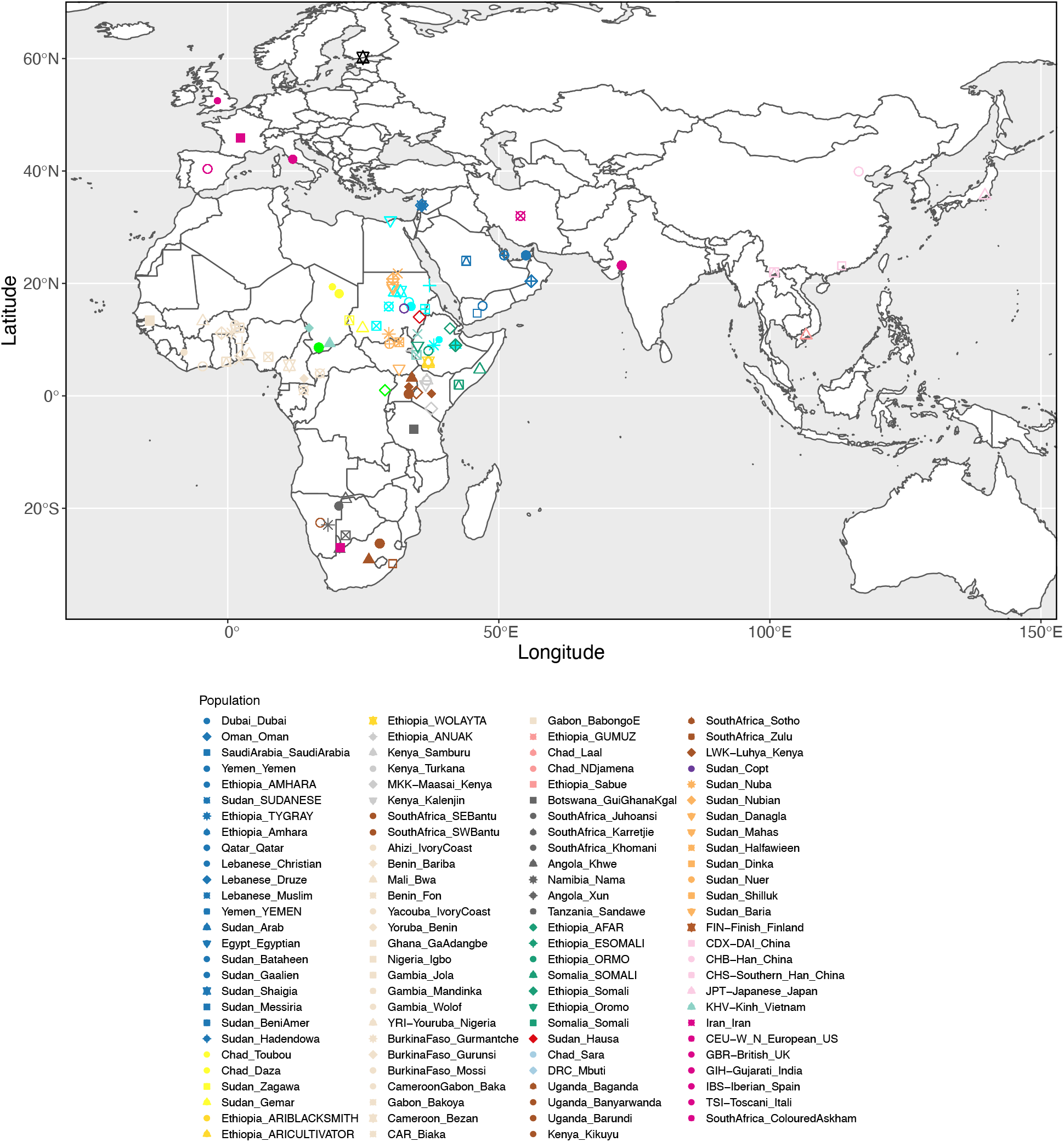
Location for populations included in this study. Colours indicates linguistic groups.

**Supplementary Figure 2:**
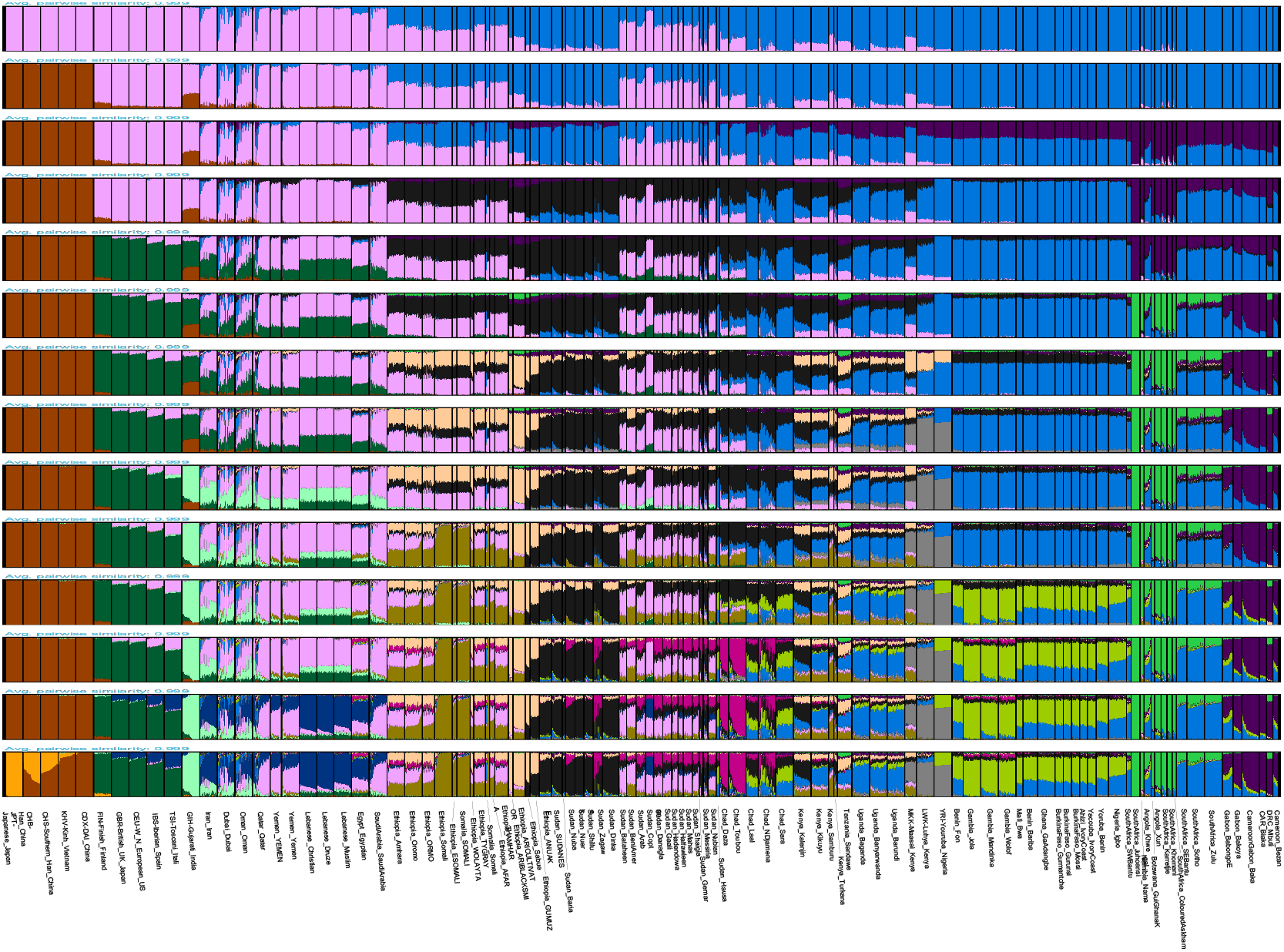
PONG visualization of 15 K’s of unsupervised ADMIXTURE analysis with 50 iterations for the full dataset. The best identified K trough cross validation was K=13

**Supplementary Figure 3:**
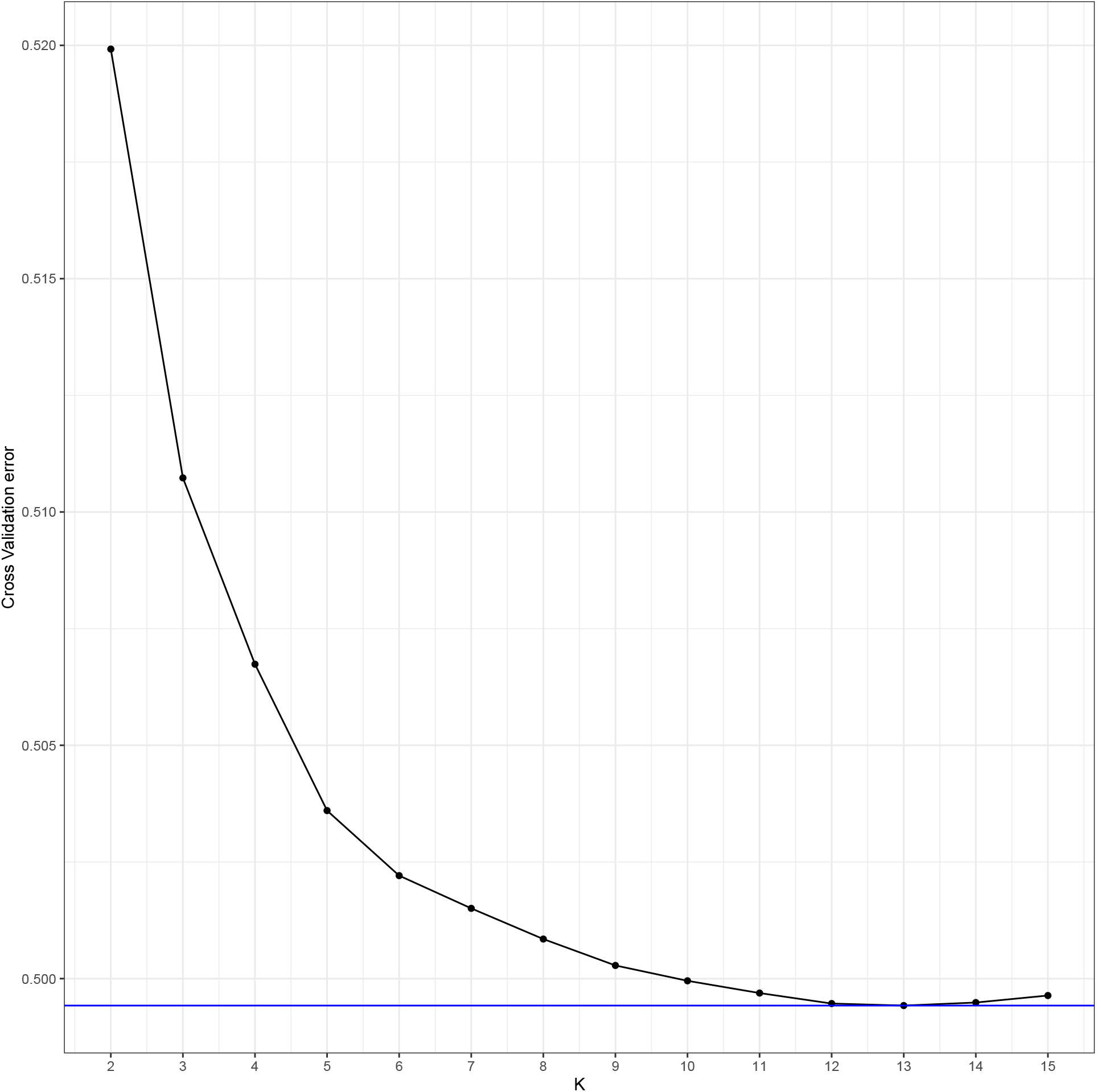
Average cross validation (CV) error for the 50 repetitions. The K with the lowest CV error was K=13, indicated by the horizontal line.

**Supplementary Figure 4:**
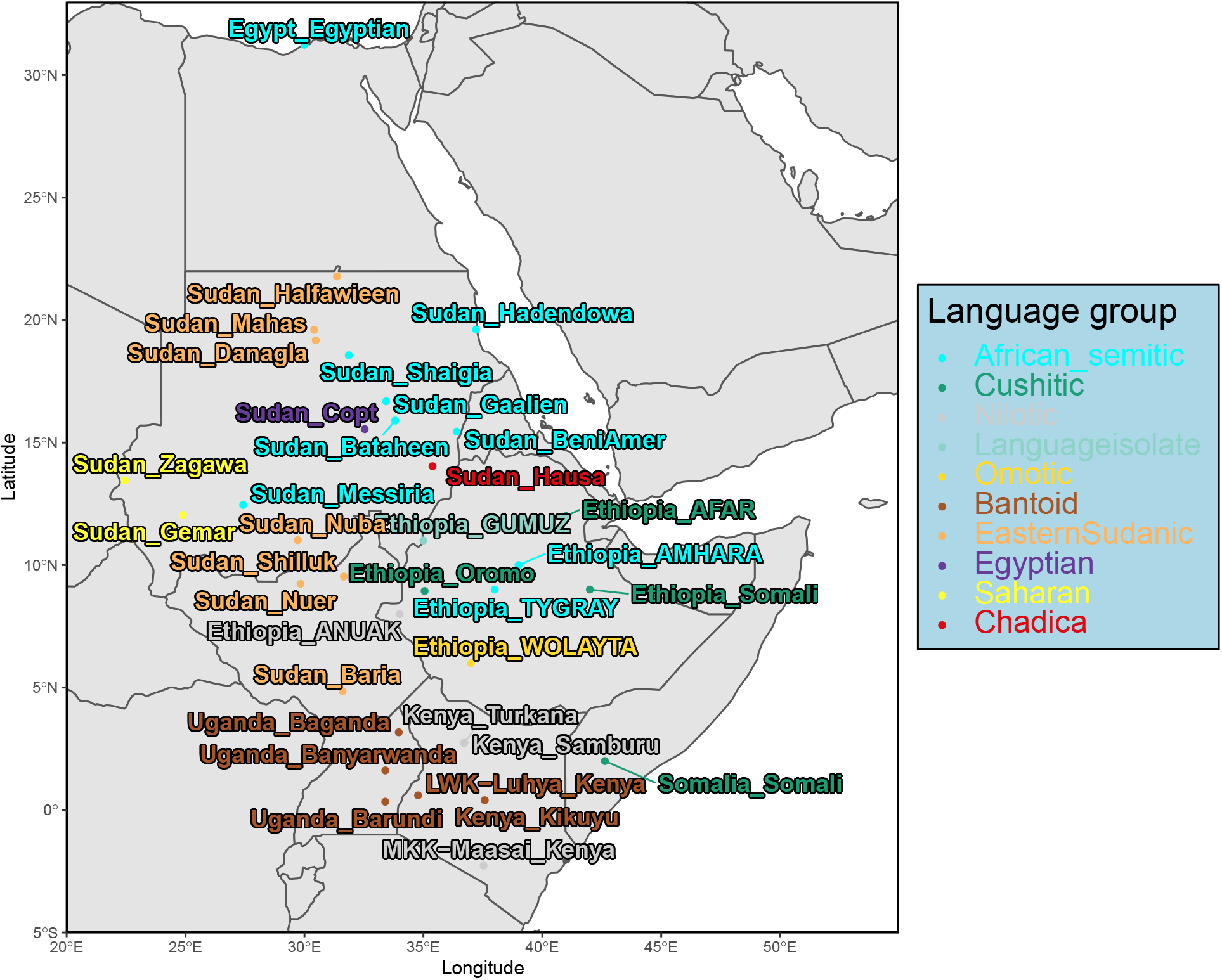
North-East African target populations, labels by country and colouring by linguistic family. African Semitic was used just more easily distinguish between the investigated populations (target) and the Middle Eastern Semitic populations.

**Supplementary Figure 5:**
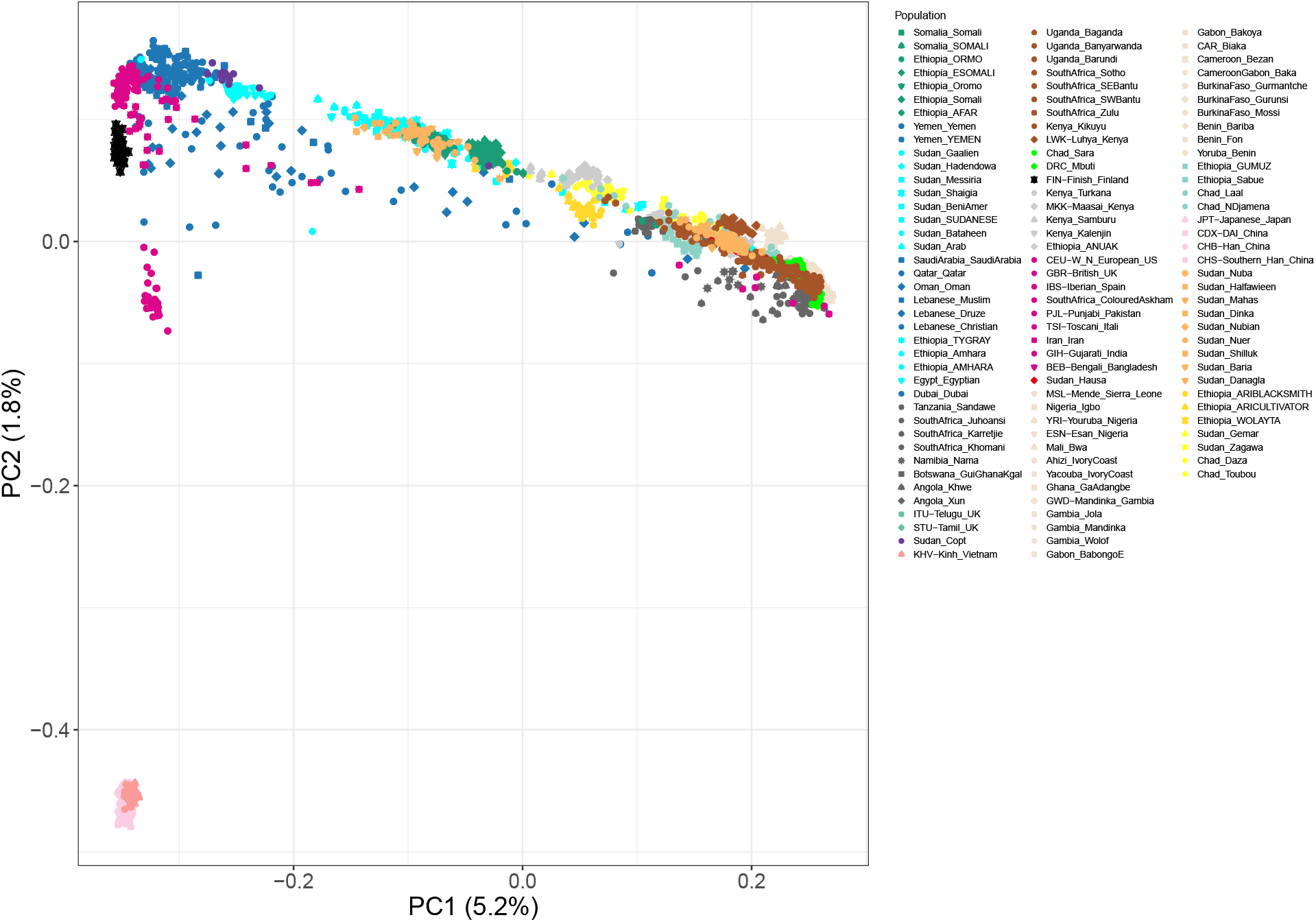
Principal Component analysis, with each value in the PCA plots is the projection of the data on the eigenvectors, scaled by the eigenvalues values within parenthesis is the PC loading. Populations are coloured by linguistic group.

**Supplementary Figure 6:**
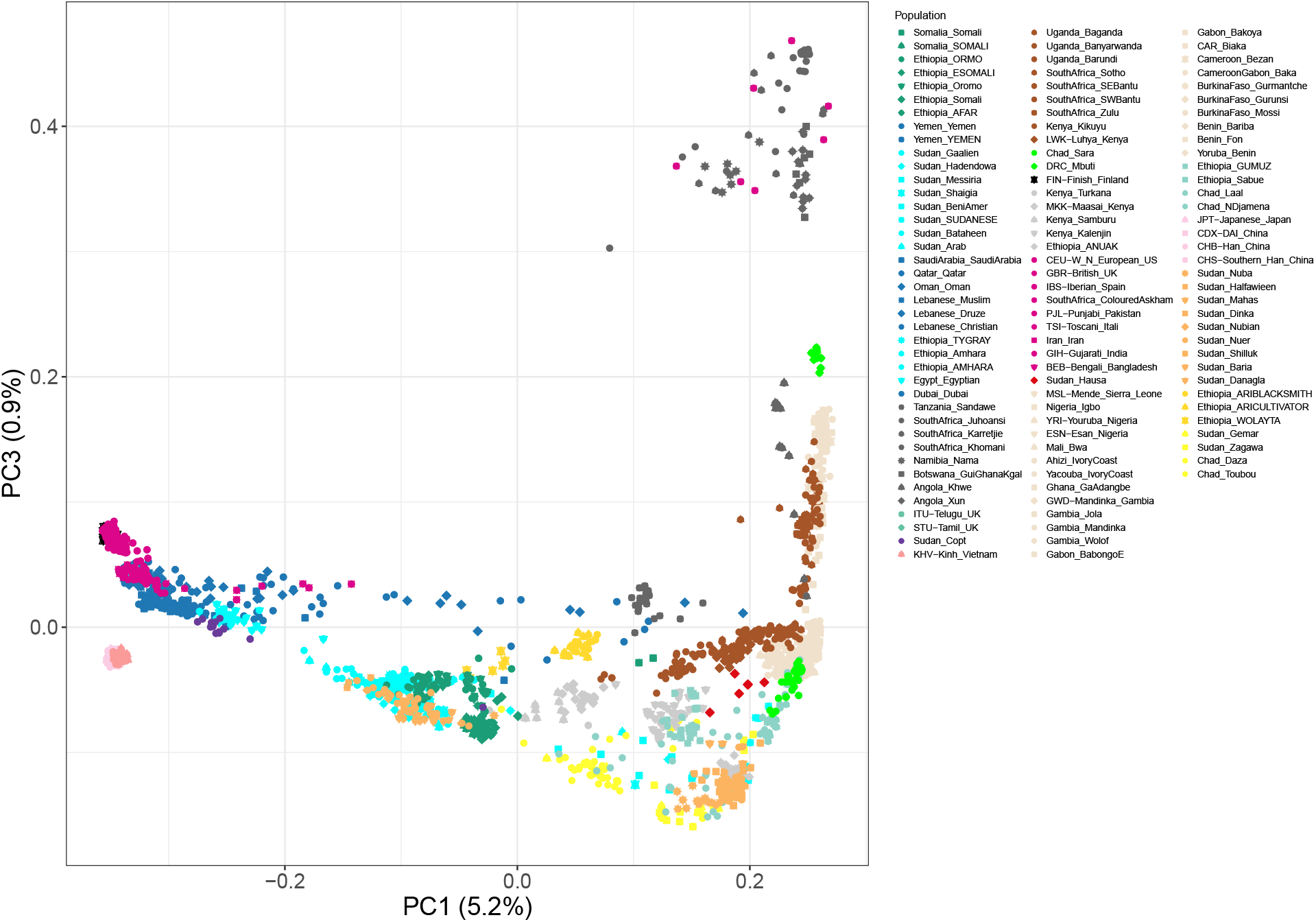
See Supplementary Figure 5 for detailed caption

**Supplementary Figure 7:**
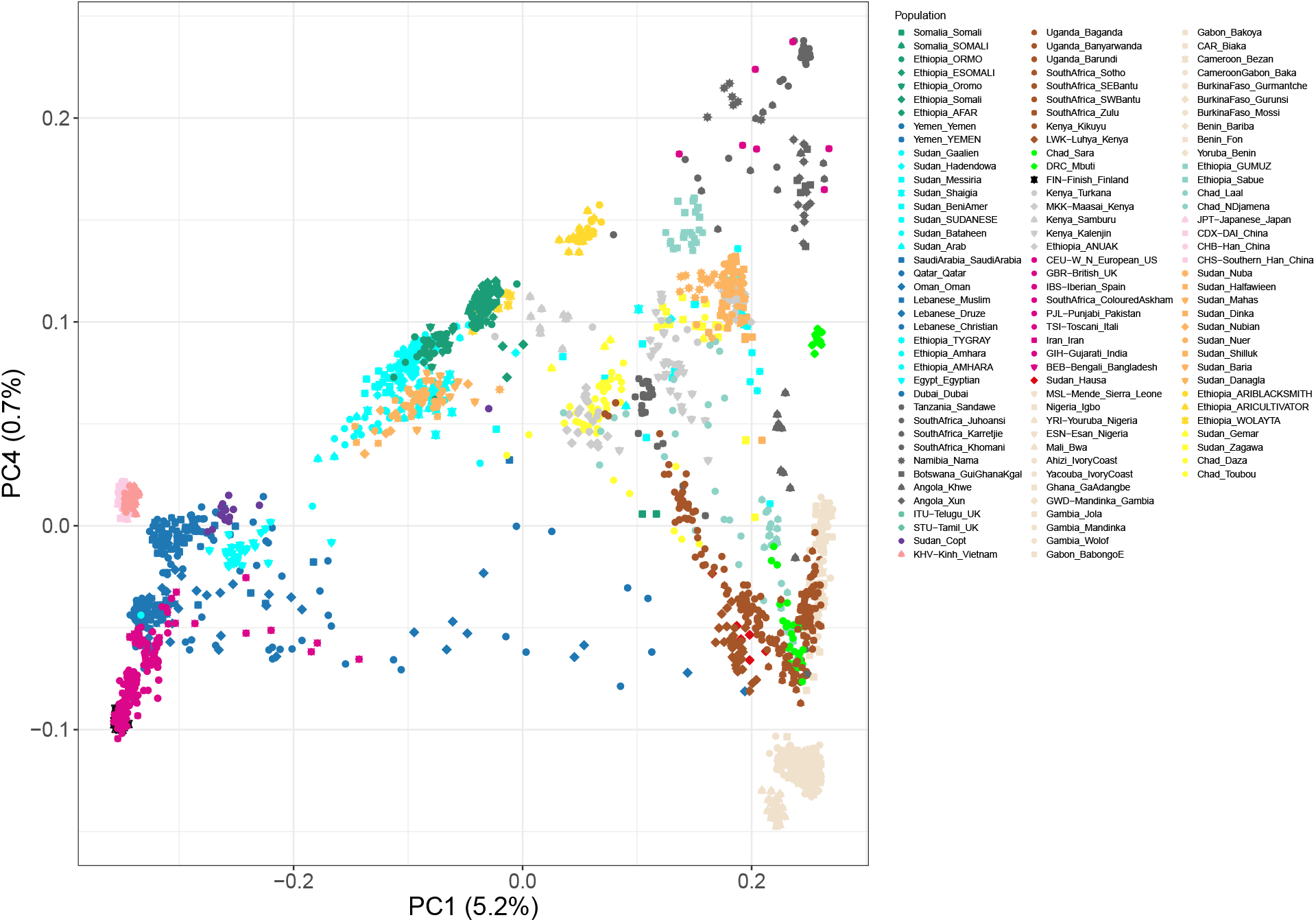
See Supplementary Figure 5 for detailed caption

**Supplementary Figure 8:**
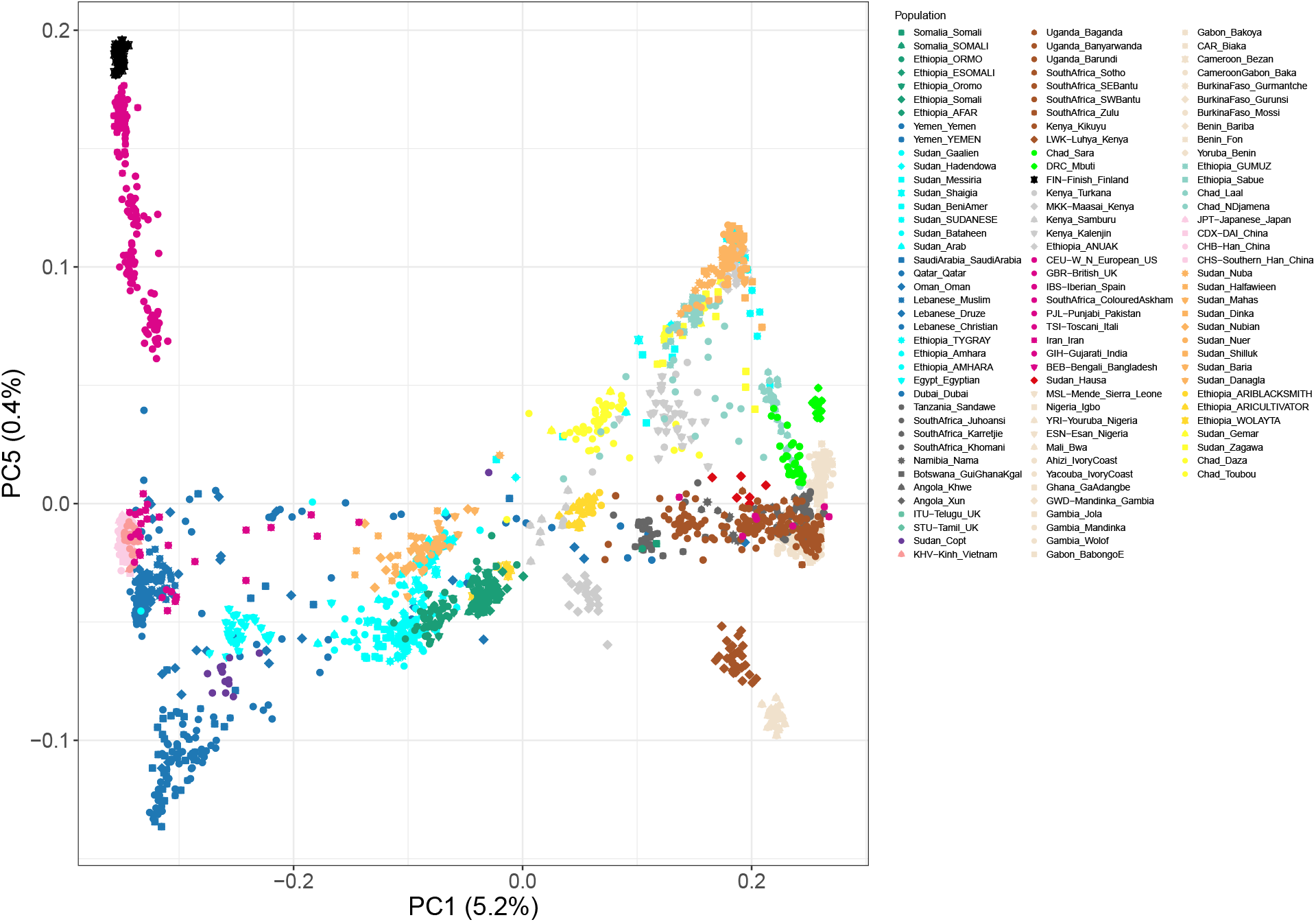
See Supplementary Figure 5 for detailed caption

**Supplementary Figure 9:**
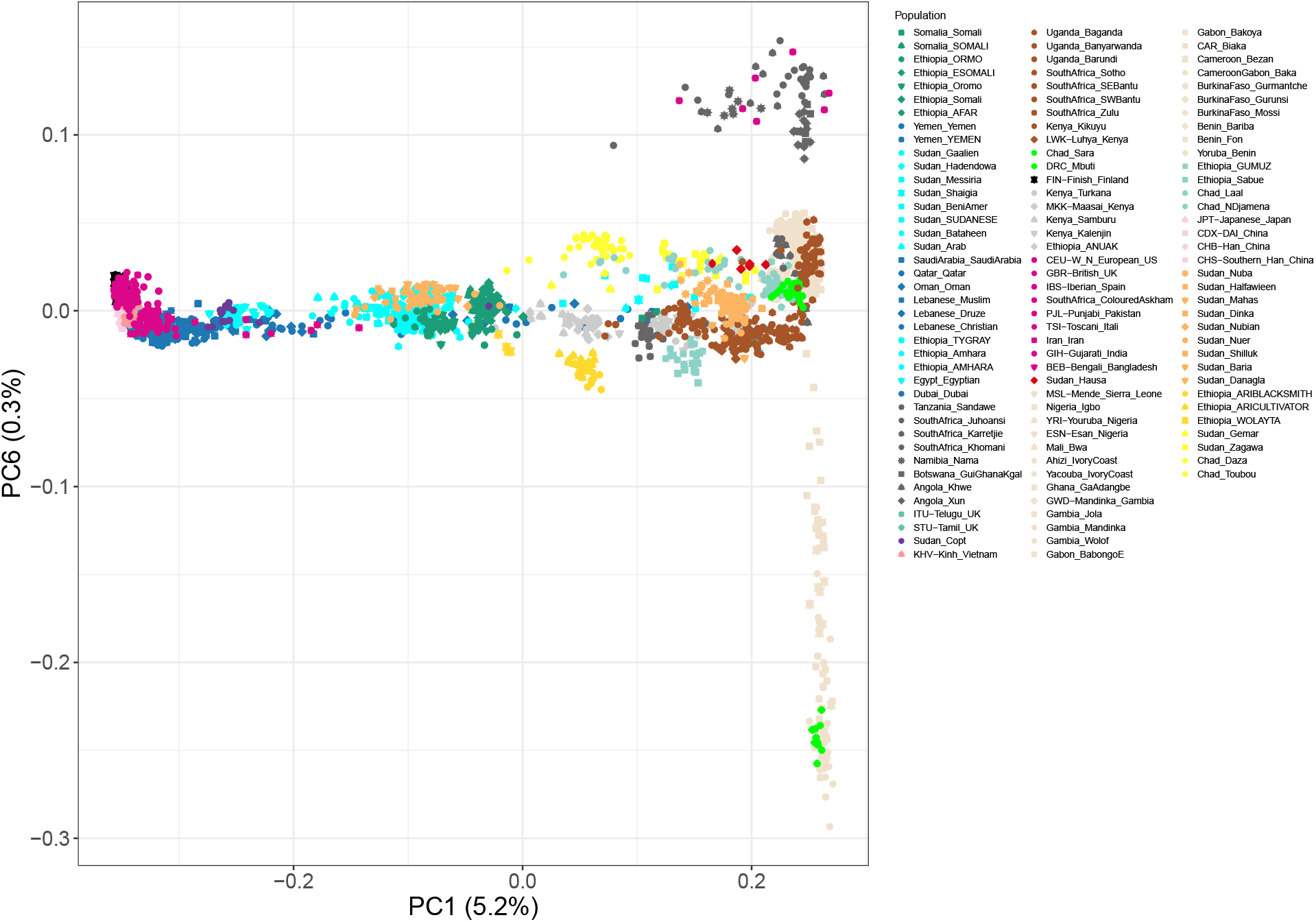
See Supplementary Figure 5 for detailed caption

**Supplementary Figure 10:**
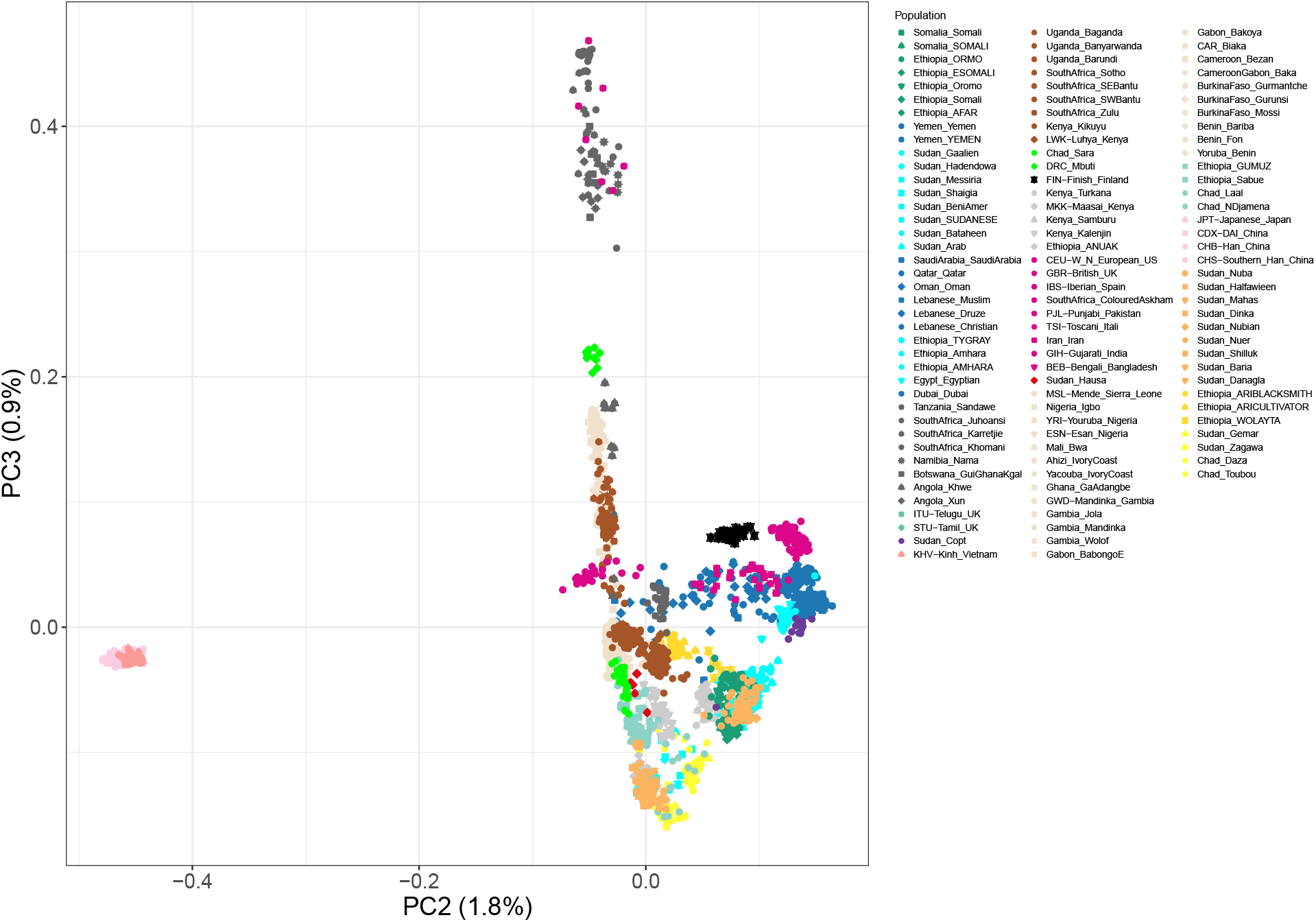
See Supplementary Figure 5 for detailed caption

**Supplementary Figure 11:**
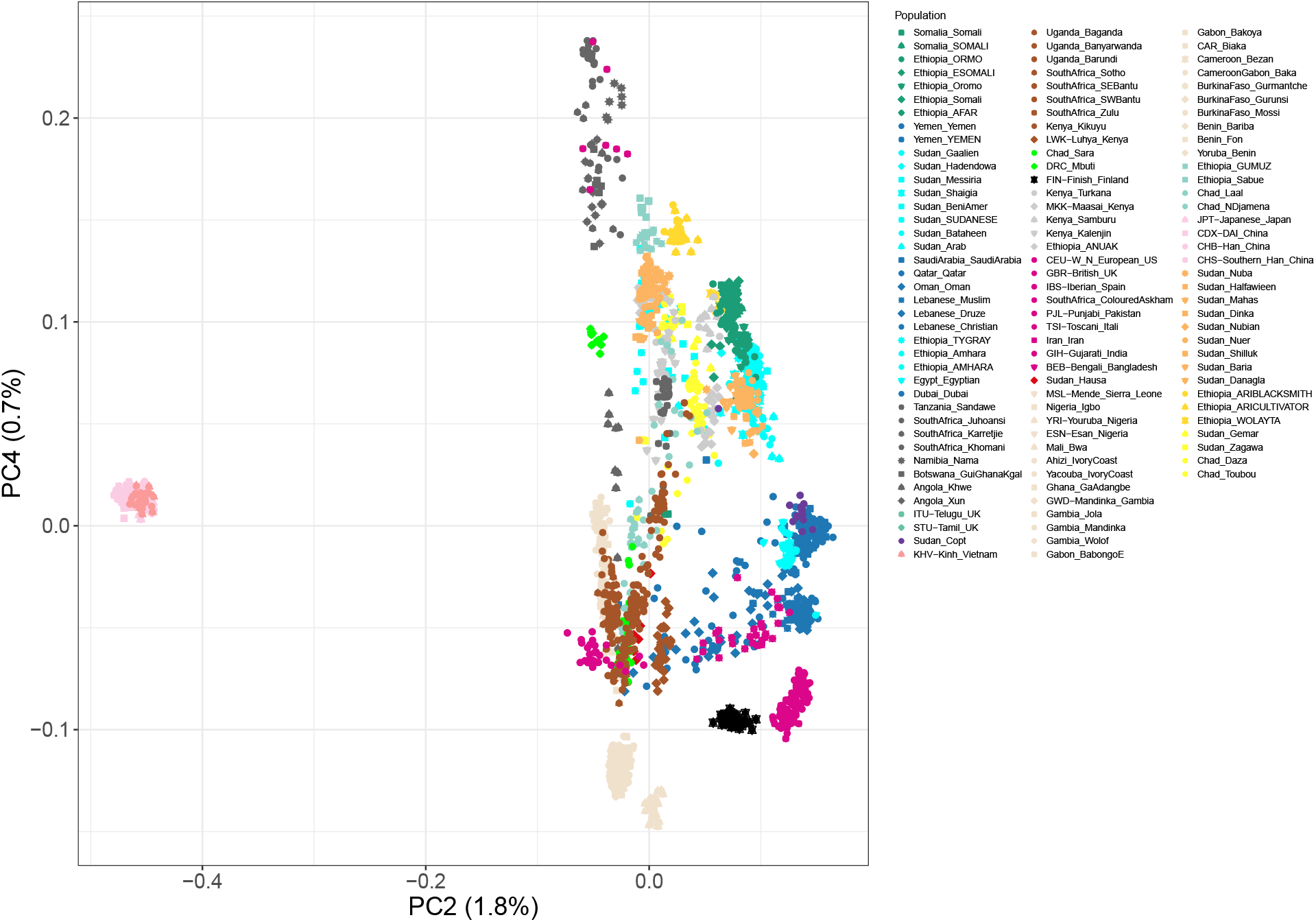
See Supplementary Figure 5 for detailed caption

**Supplementary Figure 12:**
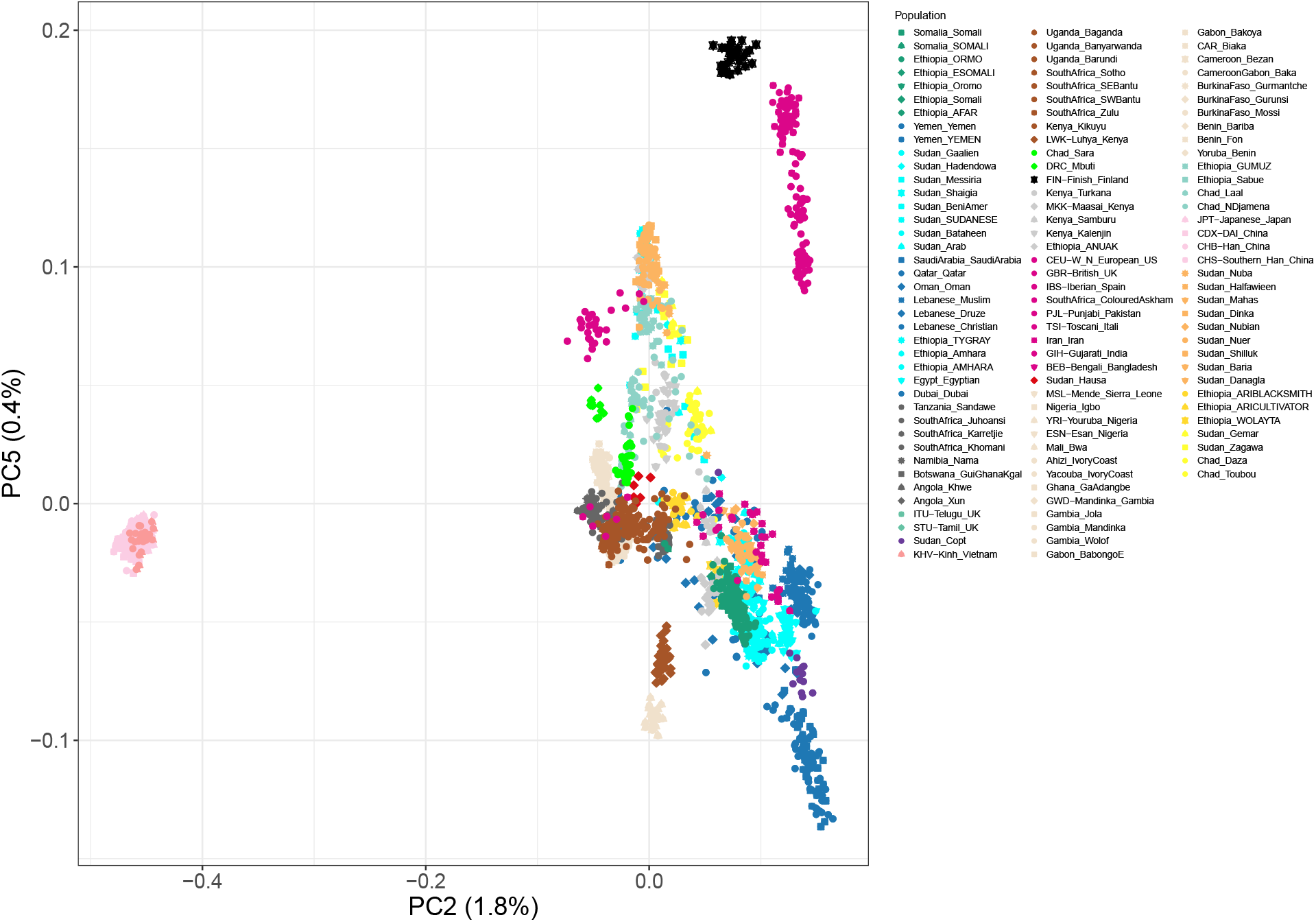
See Supplementary Figure 5 for detailed caption

**Supplementary Figure 13:**
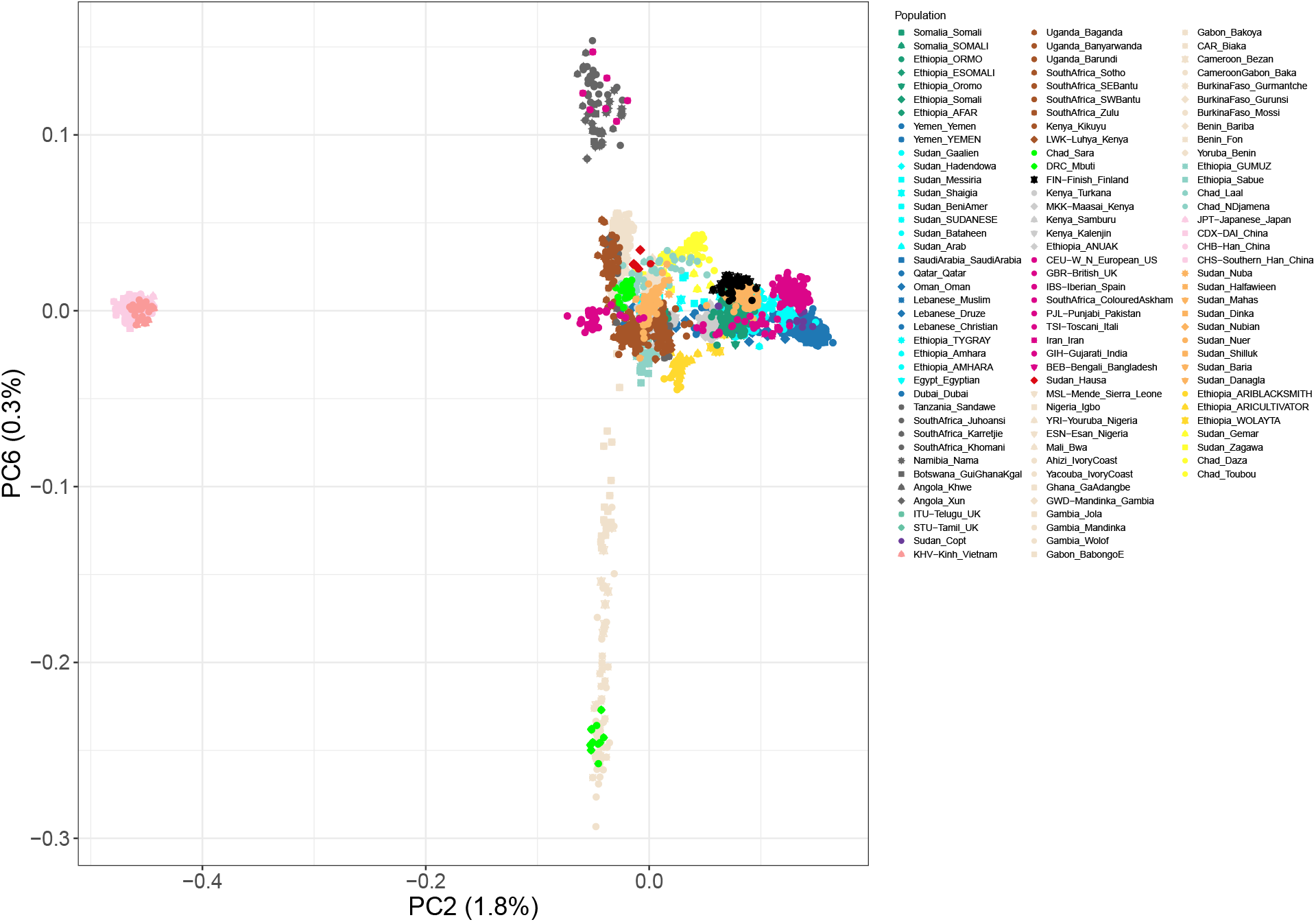
See Supplementary Figure 5 for detailed caption

**Supplementary Figure 14:**
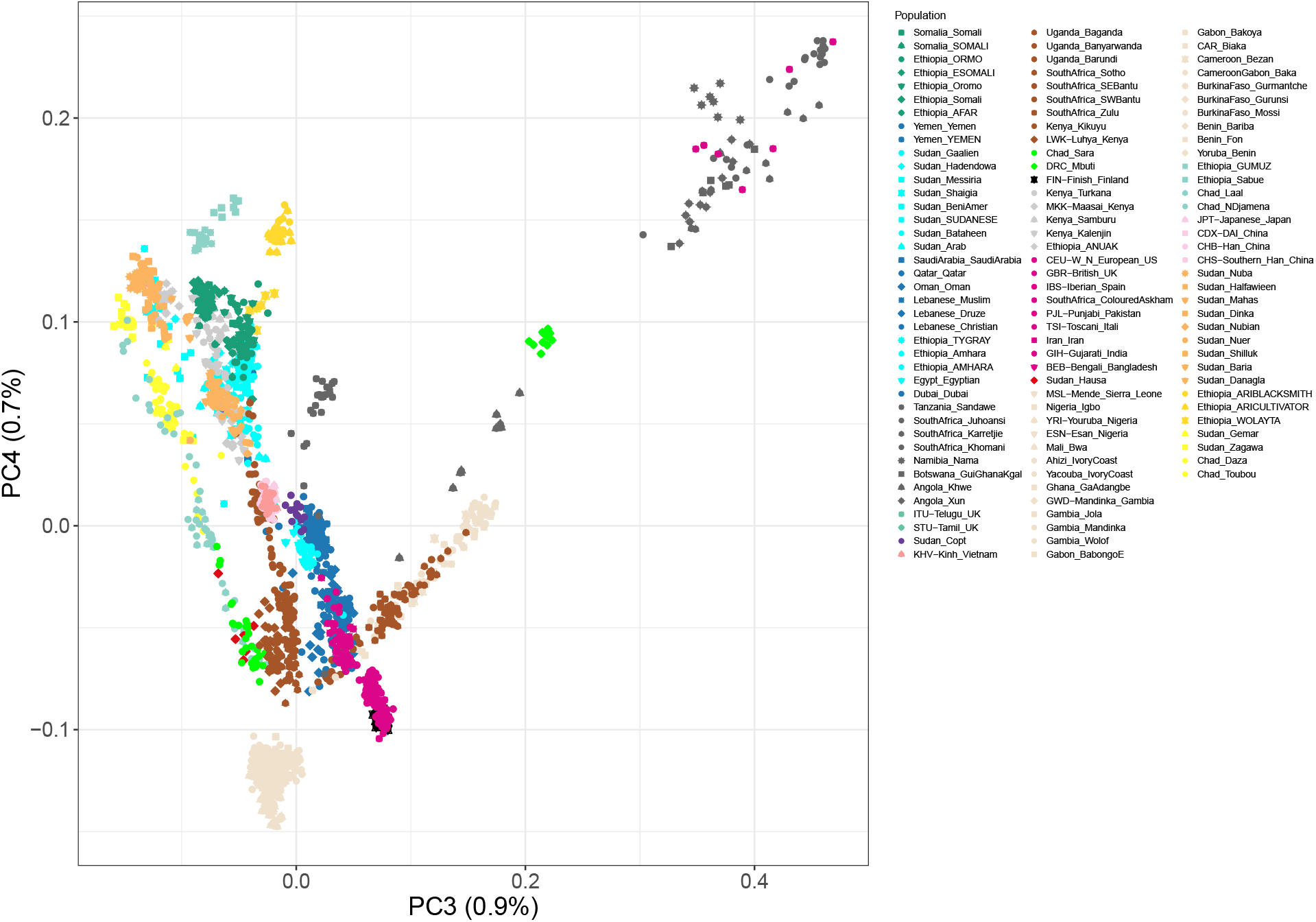
See Supplementary Figure 5 for detailed caption

**Supplementary Figure 15:**
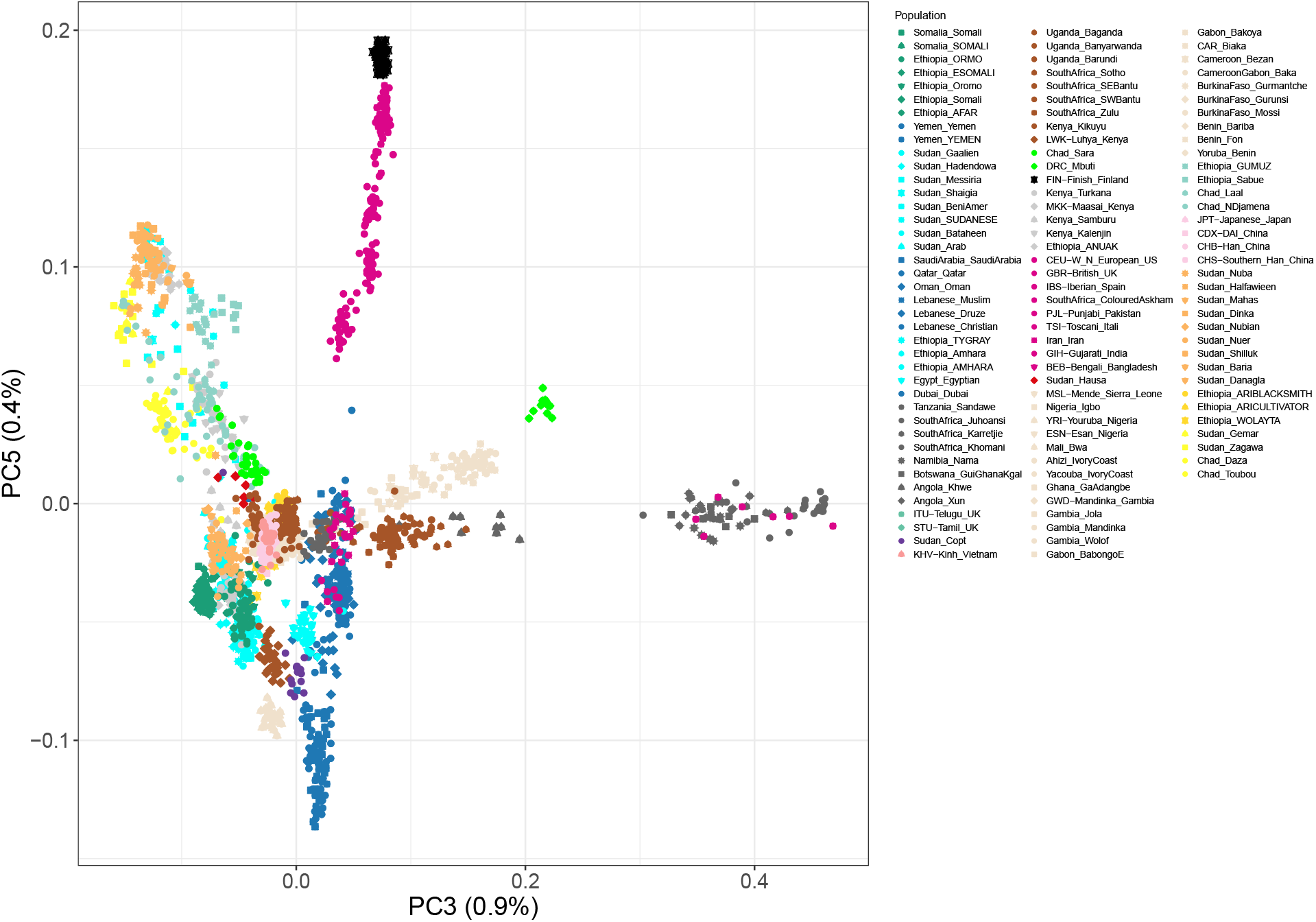
See Supplementary Figure 5 for detailed caption

**Supplementary Figure 16:**
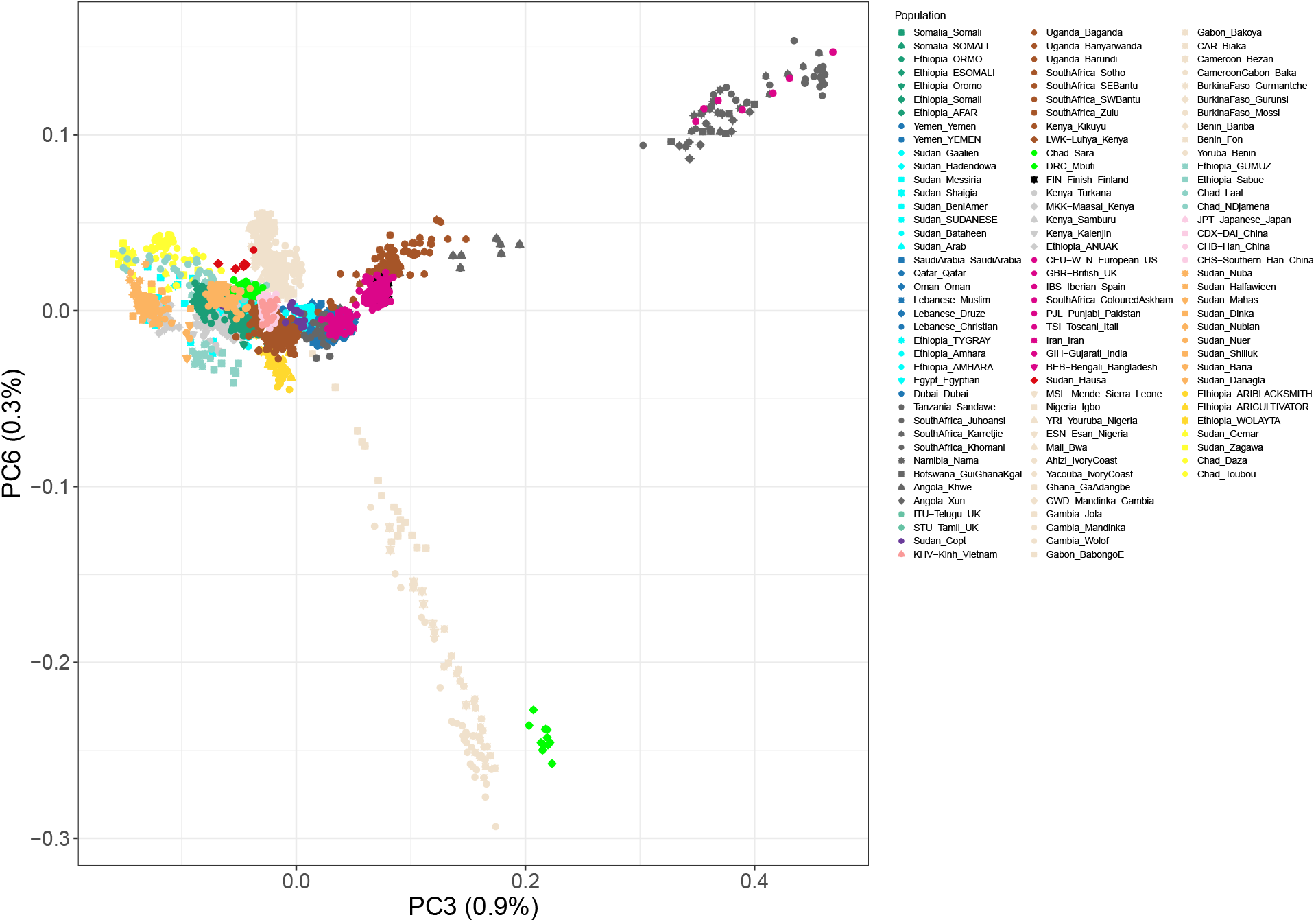
See Supplementary Figure 5 for detailed caption

**Supplementary Figure 17:**
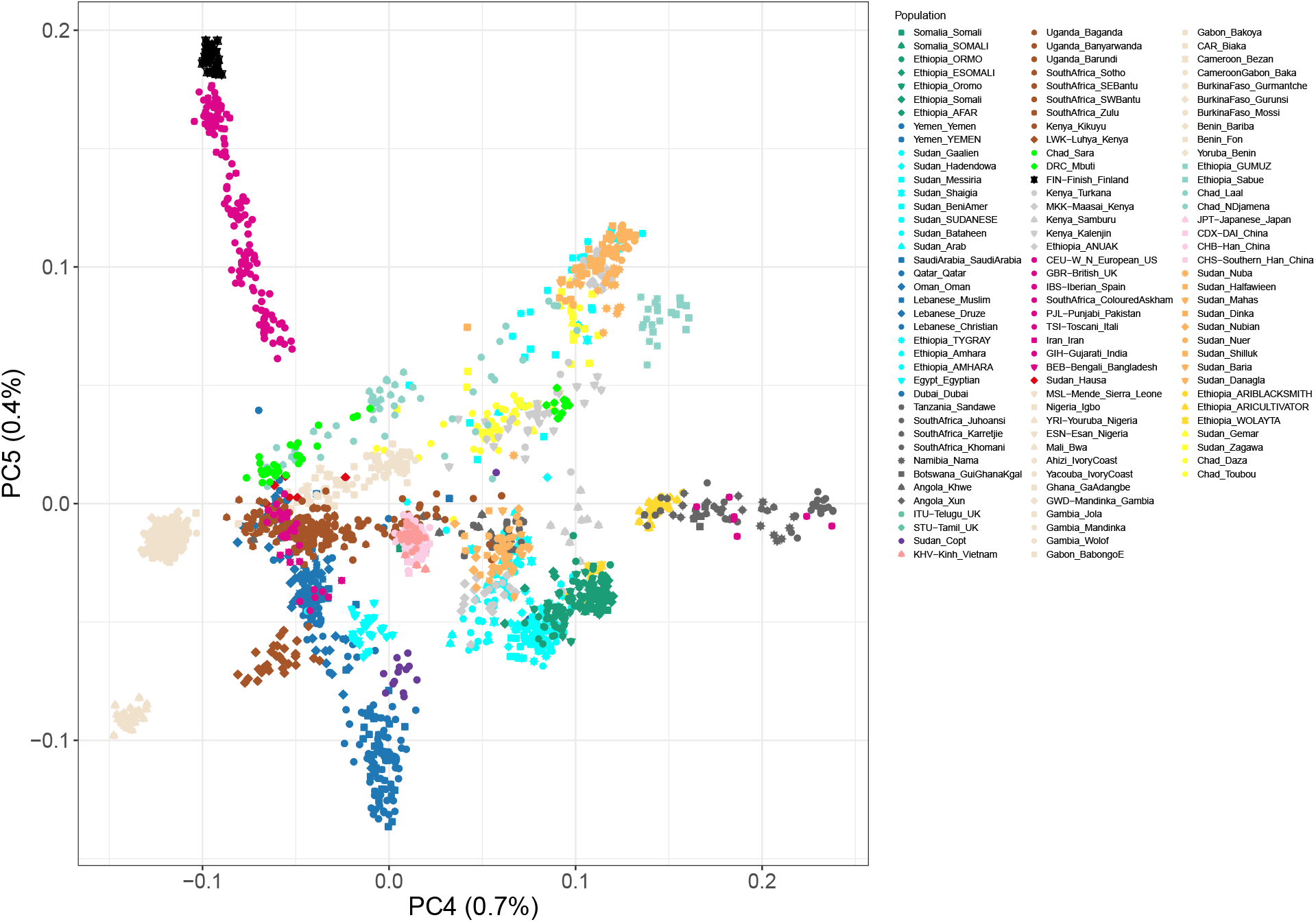
See Supplementary Figure 5 for detailed caption

**Supplementary Figure 18:**
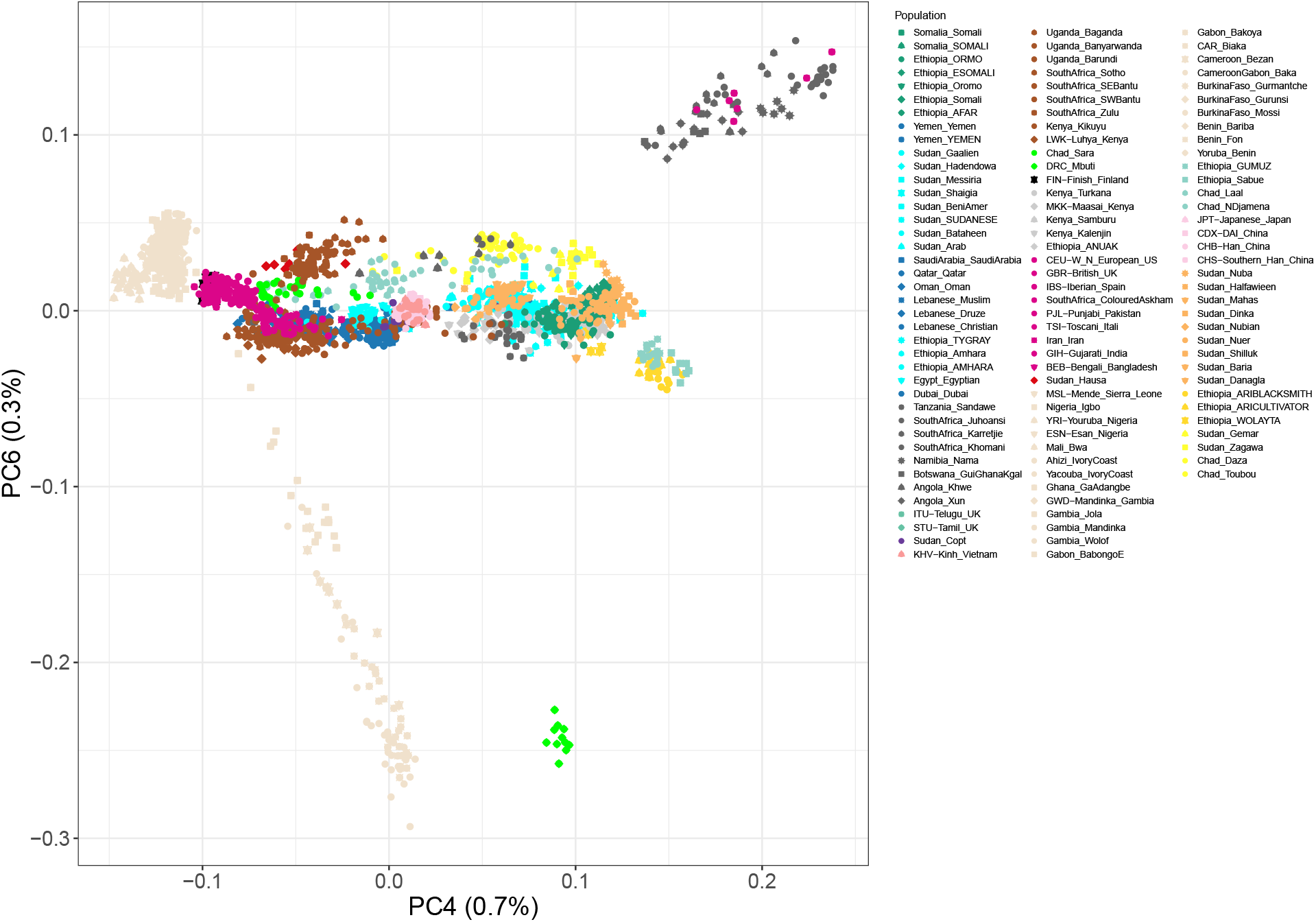
See Supplementary Figure 5 for detailed caption

**Supplementary Figure 19:**
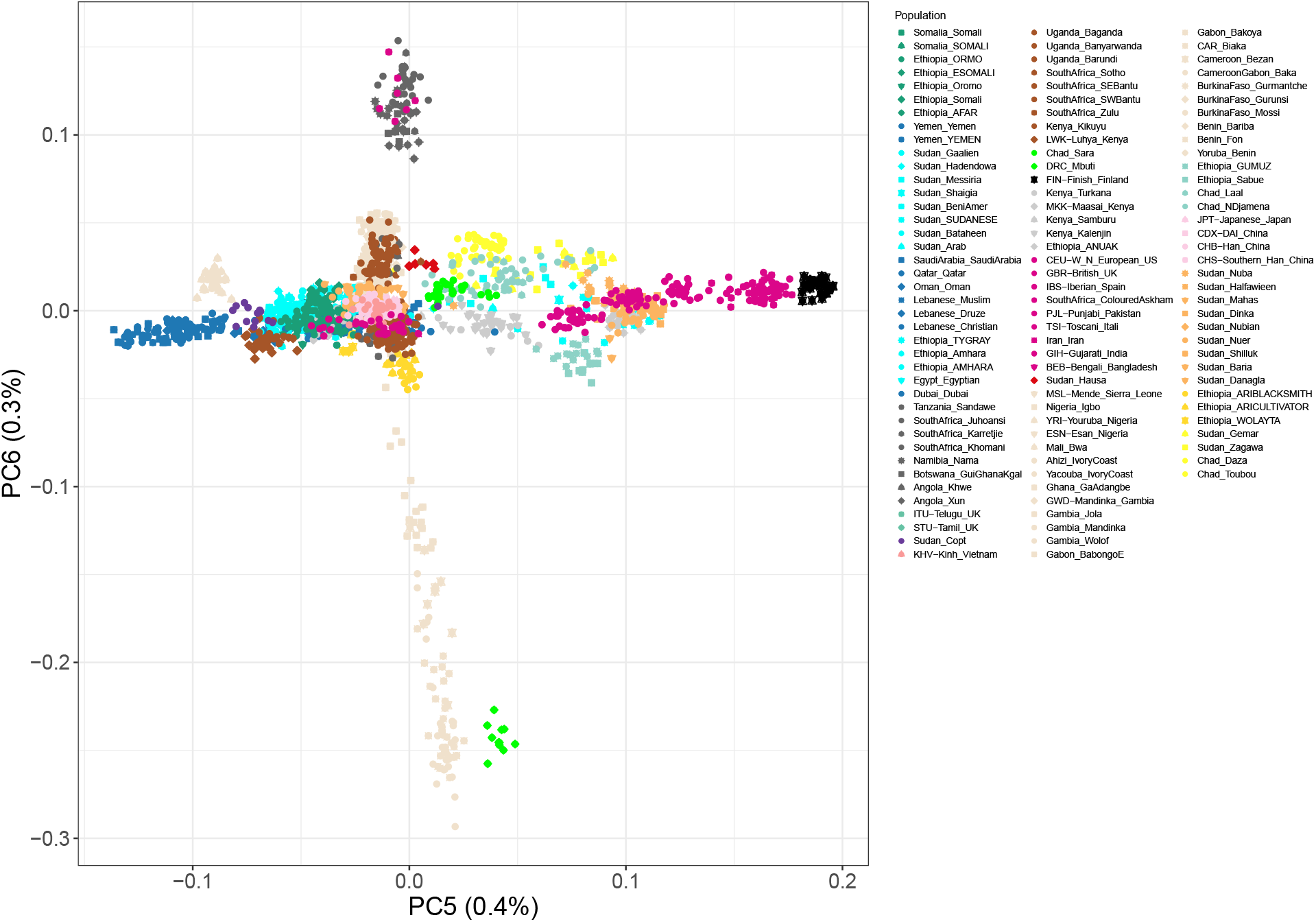
See Supplementary Figure 5 for detailed caption

**Supplementary Figure 20:**
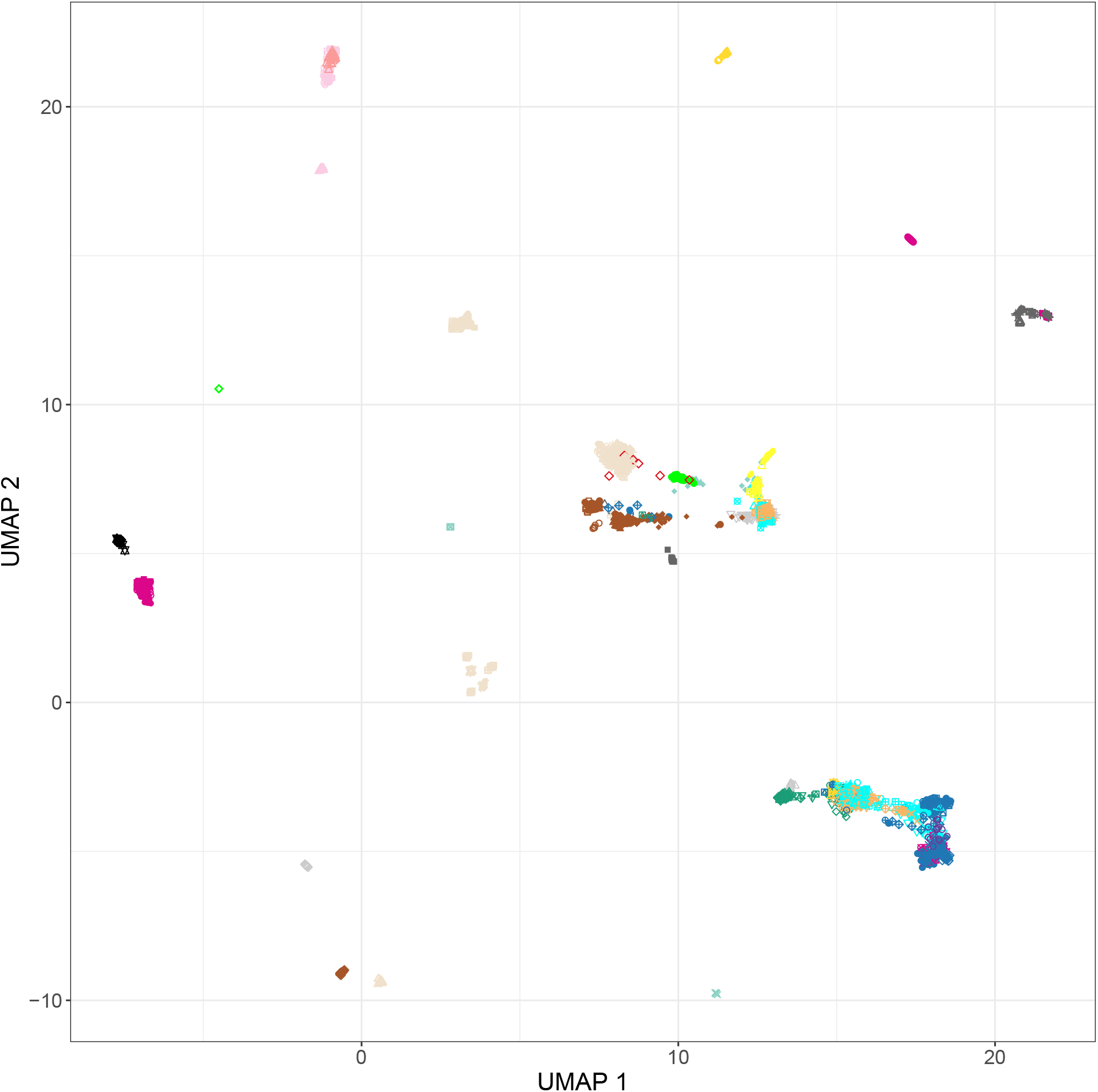
Uniform Manifold Approximation and Projection for Dimension Reduction on the full dataset, colours are the same as in 1.

**Supplementary Table 4:** Two way ANOVA statistics for the two different models, comparing the Admixture date with the factors Country, Linguistic group and Larger linguistic family (Meta.Lang.Group). The Asterisk indicates significant values. In A it’s the admixture date obtained from the best source determined by F_3_ outgroup and in B it’s the dates of admixture for the source with the highest R^2^ value.

| A | By F3 |  |  |  |  |
| --- | --- | --- | --- | --- | --- |
|  | Df | Sum_Sq | Mean_Sq | F_value | Pr(>F) |
| factor(Meta.Lang.Group) | 3 | 696.9 | 232.29 | 3.514 | 0.0352* |
| factor(Lang.Group) | 6 | 177.1 | 29.51 | 0.446 | 0.8385 |
| factor(Country) | 6 | 530.7 | 88.45 | 1.338 | 0.2890 |
| Residuals | 19 | 1255.9 | 66.10 |  |  |
| B | By R2 |  |  |  |  |
|  | Df | Sum_Sq | Mean_Sq | F_value | Pr(>F) |
| factor(Meta.Lang.Group) | 3 | 493.3 | 164.42 | 3.038 | 0.0595 |
| factor(Lang.Group) | 5 | 433.0 | 86.59 | 1.600 | 0.2166 |
| factor(Country) | 5 | 211.3 | 42.26 | 0.781 | 0.5780 |
| Residuals | 16 | 866.0 | 54.12 |  |  |

**Supplementary Figure 21:**
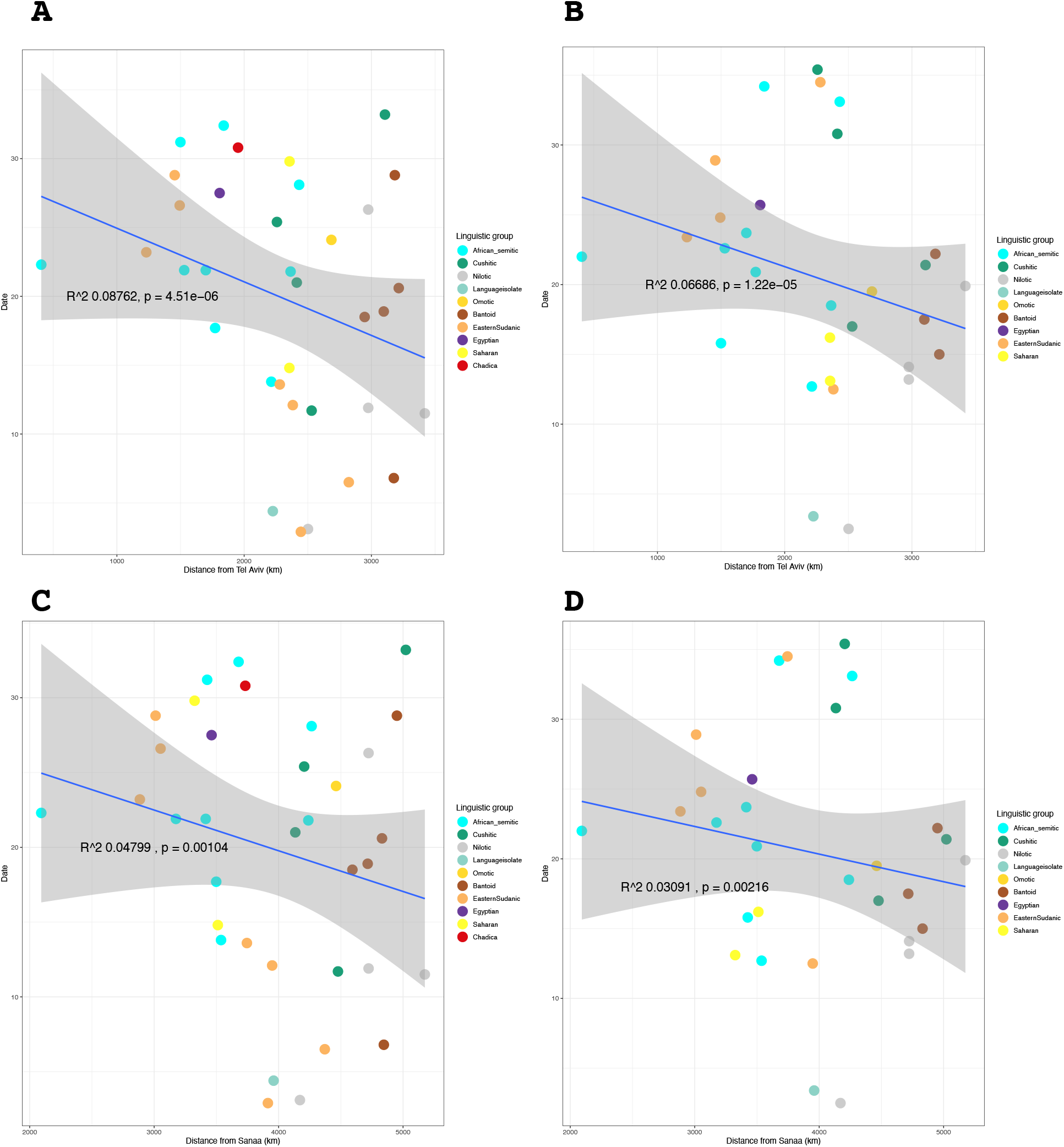
Linear regression comparing the great circle distance in kilometers from Tel Aviv and Sanaa respectively, compared to admixture date of the Eurasian ancestry estimations. Blue line is the fitted linear regression line and the grey area represent the 95% confidence interval of the standard error. A) Distance from Tel Aviv for the best by F_3_ dataset. B) Distance from Tel Aviv for the best by R^2^ dataset. C) Distance from Sanaa for the best by F_3_ dataset. D) Distance from Sanaa for the best by R^2^

